# Viral challenges and adaptations between Central Arctic Ocean and atmosphere

**DOI:** 10.1101/2024.03.17.584458

**Authors:** Janina Rahlff, George Westmeijer, Julia Weissenbach, Alfred Antson, Karin Holmfeldt

## Abstract

Aquatic viruses act as key players in shaping microbial communities. In polar environments, they face significant challenges like limited host availability and harsh conditions. However, due to restricted ecosystem accessibility, our understanding of viral diversity, abundance, adaptations, and host interactions remains limited. To fill this knowledge gap, we studied viruses from atmosphere-close aquatic ecosystems in the Central Arctic and Northern Greenland. Aquatic samples for virus-host analysis were collected from ~60 cm depth and the submillimeter surface microlayer (SML) during the Synoptic Arctic Survey 2021 on icebreaker Oden in Arctic summer. Water was sampled from a melt pond and open water before undergoing size-fractioned filtration and followed by genome-resolved metagenomic and cultivation investigations. The prokaryotic diversity in the melt pond was considerably lower compared to open water. The melt pond was dominated by a Flavobacterium sp. and *Aquiluna* sp., the latter having a relatively small genome size of 1.2 Mb and the metabolic potential to generate ATP using the phosphate acetyltransferase-acetate kinase pathway. Viral diversity on the host fraction (0.2 – 5 µm) of the melt pond was strikingly limited compared to open water. From 1154 dereplicated viral operational taxonomic units (vOTUs), of which two-thirds were predicted bacteriophages, 17.2% encoded for auxiliary metabolic genes (AMGs) with metabolic functions. Some AMGs like glycerol-3-phosphate cytidylyltransferase and ice-binding like proteins might serve cryoprotection of the host. Prophages were often associated with SML genomes, and two active prophages of a new viral genera from the Arctic SML strain *Leeuwenhoekiella aequorea* Arc30 were induced. We found evidence that vOTU abundance in the SML compared to ~60 cm depth was more positively correlated to the distribution of a vOTU across five different Arctic stations. The results indicate that viruses employ elaborated strategies to endure in extreme and host-limited environments. Moreover, our observations suggest that the immediate air-sea interface serves as a platform for viral distribution in the Central Arctic.

## Background

The Central Arctic Ocean is a remote and inhospitable region located in the Arctic Circle, encompassing the waters surrounding the North Pole. The region is characterized by harsh environmental conditions including cold temperatures, storms, ice and snow coverage, and extended periods of darkness (polar night) and light (midnight sun) in Arctic winter and summer, respectively. Sea ice covers most of the area throughout the year posing significant challenges for scientific research and exploration. Sea ice coverage regulates the amount of sunlight reaching the surface of the ocean, consequently influencing primary productivity [1]. Since the Arctic region experiences rapid environmental changes due to climate change [2] understanding the Central Arctic and its ecosystems becomes increasingly important. With forecasted globally rising temperatures, more sea ice will melt and net primary productivity will increase further [3, 4]. Viruses exert top-down control by infecting microorganisms, and thus play a crucial role in aquatic ecosystems by influencing nutrient cycling and shaping microbial diversity [5, 6]. In general, viruses are understudied components of the Arctic Ocean and especially the Central Arctic Ocean (reviewed by [7]). This applies also to viruses in the upper 1 m of the ocean’s surface including the < 1 mm surface microlayer (SML), where so-called neuston organisms reside [8, 9]. The lack of microbial and viral surveys from the immediate air-sea interface can be attributed to the fact that the typical CTD Niskin rosette water samplers cannot collect SML. Especially the upper millimeter and meters of the Arctic Ocean are affected by melting ice, exerting strong salinity gradients, and supplying the oceanic water with diverse sea-ice associated biota [10].

When sea ice melts, especially thin first-year ice, microbial biopolymers consisting of proteinaceous material can contribute to the formation of a gelatinous SML by accumulating at the air-sea boundary [11]. Polymeric gels at the air-water interface were previously detected in melt ponds [11], which are pools of melted water that form on the surface of sea ice, where they can make up ~50% of the surface area [12]. Viruses of bacteria (bacteriophages) have previously been isolated from Arctic sea ice [13, 14] but the fate of viruses in melt ponds remains unknown [10]. However, by accumulating in the skin layer between ocean and atmosphere, the polymeric compounds and viruses can more easily end up in aerosols [15, 16] with the potential of nucleating ice and influencing cloud formation [17–19].

The Tara Oceans Global Circle expedition has increased our knowledge of viral populations around the Arctic Ocean [20], however, investigations on viruses from the Central Arctic Ocean are missing. This especially includes omics-based investigations of the virioneuston (reviewed by [21]), which are lacking for the whole polar regions. Virioneuston compared to virioplankton activity and abundance in the Arctic (Norwegian Sea and North of Svalbard) were found to be enhanced [22], suggesting that viral lysis in Arctic SML, compared to other ecosystems like sea ice [23], can be an important factor in the release of dissolved organic carbon promoting growth of heterotrophic microbes. Therefore it fits well that also bacterioneuston numbers were higher in Arctic Ocean SML compared to underlying water [24], and the SML of open water contained more bacteria that the SML of melt ponds [11]. With increasing warming of the surface ocean under ongoing climate change, bacterial production is enhanced and triggers viral activity [25], resulting from the tight coupling of viral and bacterial variations in the Nordic Seas, which get shaped by different abiotic factors like water masses, latitude, and water pH [26]. During the Synoptic Arctic Survey 2021 expedition, we aimed to fill major knowledge gaps regarding diversity, lifestyle prevalence, host interactions and dispersal patterns of DNA viruses by conducting samplings of the SML during our journey through the Central Arctic. We used metagenomics, cultivation, and prophage induction assays to gain insights into how Arctic viruses adapt to hosts in the boundary layer ecosystem influenced by freeze-thaw cycles.

## Methods

### Sampling and water filtration

Samples were collected during the Synoptic Arctic Survey 2021 [27] exploring the Central Arctic Ocean, the North Pole, and Northern Greenland between July and September 2021 (see Table S1). Wind speed was measured at the sampling site using a handheld anemometer model MS6252A (Mastech Group, Brea, CA, USA). Water temperature and salinity were measured using a thermosalinometer, model Professional Plus, YSI (Xylem, Washington, D.C., USA). Sampling of SML was conducted with a glass plate sampler [28] either from a) a melt pond, b) seawater sampled from the ice edge of a lead (Figure 1b), and c) from seawater reached from the lowered gangway of the icebreaker Oden. The glass plate was submerged perpendicularly to the water surface into the water and slowly withdrawn. Thereby SML adheres to the glass and is collected from it by wiping off both sides of the plate with a squeegee blade into a collection bottle. Glass plate and bottle were cleaned with ethanol and pre-rinsed with sample water. This method is state-of-the-art for virioneuston sampling [29, 30].

**Figure 1:**
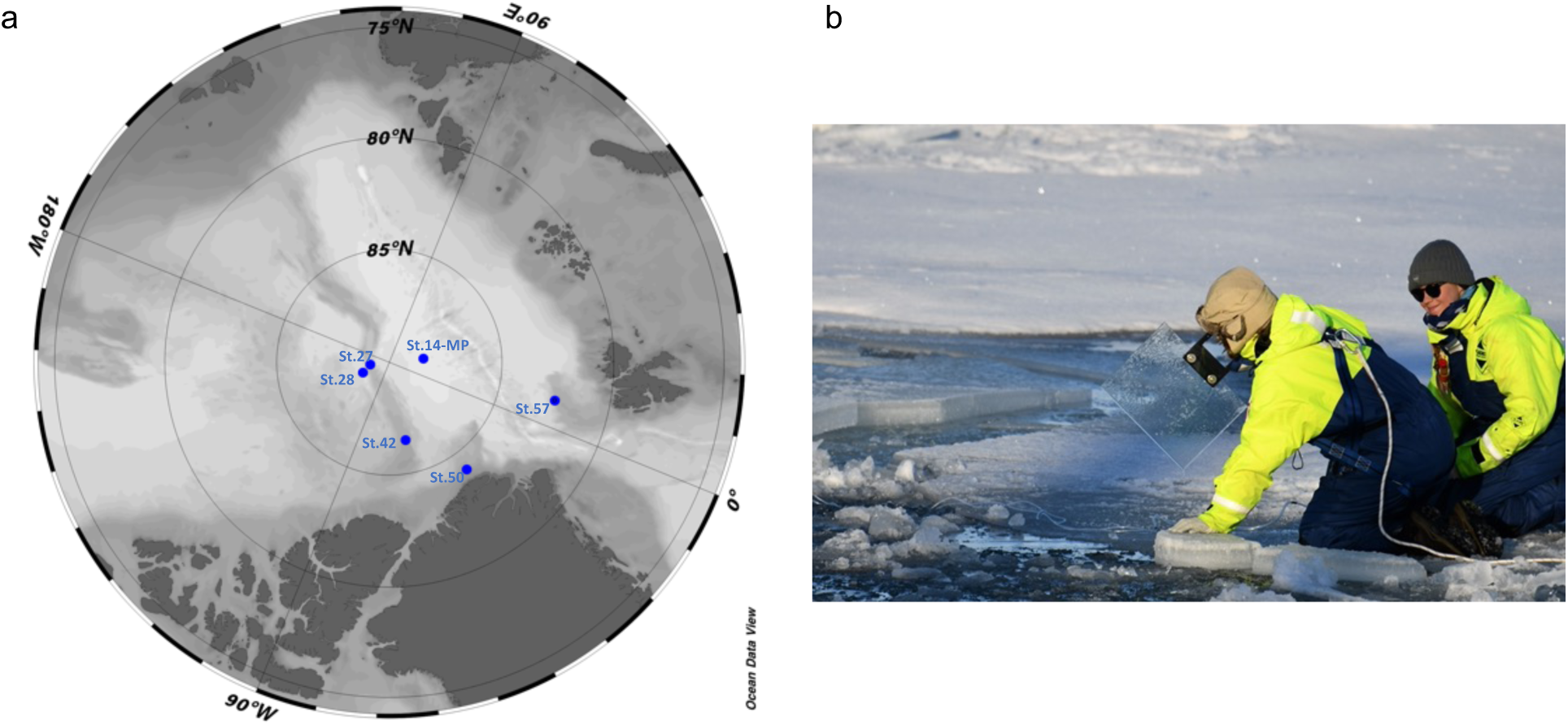
Sampling the sea-surface microlayer in the Central Arctic. Map showing stations of surface microlayer and underlying water sampling. (a) “-MP”: melt pond station. Map was created with Ocean Data View [33]. (b) Representative picture for sampling surface microlayer with the glass plate sampler from the ice edge (Photo: Hans-Jørgen Hansen, MacArtney Underwater Technology).

Sampling locations for the sporadic microlayer samplings are depicted in Figure 1a. As a reference sample, subsurface water (SSW) from ~60 cm depth was sampled using a 100 ml syringe connected to a hose. The syringe and the hose were rinsed with sample water and sterilized with 70 % ethanol and MilliQ water after use. Sample water was sequentially vacuum-filtered over a series of membranes featuring pore sizes of 5 µm and 0.2 µm (47 mm diameter, both Nuclepore^TM^ track-etched polycarbonate membrane, Whatman, Maidstone, UK) and the flow-through was flocculated using iron-III-chloride (FeCl_3_) solution [31]. A 10 x higher concentration than in the original protocol of 10 mg FeCl_3_ L^−1^ was chosen to recover more viruses as this concentration was recently recommended for freshwater [32], and SML samples were partly derived from freshwater (a melt pond) or affected by freshwater melting ice at the air-water boundary. The iron flocculates were filtered with a peristaltic pump onto a 142 mm, 1 µm pore-sized polycarbonate filters (Whatman). From station 42 onwards, we had to switch to a 1 µm PTFE membrane (Omnipore, Merck Millipore, Darmstadt, Germany). Filters were stored on dry ice or −80 °C until DNA extraction.

### DNA extraction and metagenomic analysis of MAGs

DNA was extracted using DNAeasy Power Soil Pro kit or DNAeasy PowerMax Soil kit (Qiagen, Kista, Sweden) for 47 mm and 142 mm filters, respectively. When applying the PowerMax Soil kit, a subsequent ethanol precipitation step with glycogen from mussels (Roche Diagnostics, Basel, Switzerland) as carrier molecule was performed. DNA concentration was measured on a Qubit^®^ 2.0 Fluorometer (Invitrogen/ Life Technologies Corporation, Carlsbad, CA, USA). Short-read sequencing was performed by SciLifeLab (Solna, Sweden) using Illumina DNA PCR-free library preparation. Samples were sequenced on NovaSeq6000 (NovaSeq Control Software 1.7.5/RTA v3.4.4) with a 151nt (Read1)-19nt (Index1)-10nt (Index2)-151nt (Read2) setup using ‘NovaSeqXp’ workflow in ‘S4’ mode flowcell. The Bcl to FastQ conversion was performed using bcl2fastq_v2.20.0.422 from the CASAVA software suite. The quality scale used is Sanger / phred33 / Illumina 1.8+. Adapter trimming, and quality control were conducted using bbduk as part of BBTools [34] under application of contaminant filtering using the Illumina PhiX spike-in reference genome (phix174_ill.ref.fa) and the artificial contaminants file (sequencing_artifacts.fa). Sickle v.1.33 [35] was subsequently run in pe mode and -t sanger setting. mOTUs v.3.0.2 [36, 37] was used on trimmed reads with options -A (reports full taxonomy) -c (reports counts) -M (to save intermediate marker gene cluster count) for the taxonomic profiling of bacteria and archaea based on single-copy phylogenetic marker genes. Data were read sum normalized and analyzed in R v.4.0.3 [38] within R Studio using the packages phyloseq v.1.34.0 [39] and ggplot2 v.3.4.2 [40]. Binning of prokaryotic metagenome-assembled genomes (MAGs) from > 5 µm and 0.2 - 5 µm samples was conducted using CONCOCT v.1.1.0 [41] and MetaBAT v.2.12.1 [42] on MetaSPAdes v.3.15.3 [43] - derived scaffolds with a minimum length of 1000 bp. DAS_Tool v.1.1.3 [44] with default threshold was used to obtain a dereplicated set of MAGs, and uBin v.0.9.14 [45] was used for subsequent manual refinement. MAGs underwent quality checks in CheckM2 v.0.1.3 [46], and their GC content, coding density, and genome size was obtained from CheckM v.1.1.3 [47]. Then taxonomic assignment was performed by the classify_wf in GTDB-Tk v.2.1.0 [48] with database R207_v2. MAGs with estimated completeness and contamination scores of ≥ 70 % and ≤ 10 %, respectively, in either uBin or CheckM2 were considered for further analysis and dereplicated together with the isolate genomes (see below) in dRep v.3.4.0 [49] using default settings. Reads were mapped back to dereplicated MAGs and isolate genomes by using the --reorder flag in Bowtie v.2.4.5 to subsequently predict *in situ* replication rates at default thresholds in iRep v.1.1.7 [50] including a mismatch filtering step with a 2 % error rate (-mm 3). To assess encoded metabolic potential of MAGs, eggnog-mapper v.2.1.9 [51] with options --itype genome -m diamond --genepred prodigal was run on MAGs and bacterial isolate genomes (see below). To determine pathway completeness, the obtained functional annotation from the eggnog-mapper was used as input for the reconstruct tool as part of the Kyoto Encyclopedia of Genes and Genomes (KEGG) mapper. Read breadth on MAGs was used to decide if a MAG is present in the melt pond or and open water station, namely if 90% of the genome had a coverage of at least 1 in at least one of the melt pond or open water stations. This was done after mapping with Bowtie 2 and mismatch (-m 3) filtration using mapped.py (https://github.com/christophertbrown/bioscripts/blob/master/ctbBio/mapped.py). We excluded potential contaminant bacteria from MAGs and isolates (see discussion in the supplement) from this analysis (labelled with a “C” in Table S2).

### Mining of vOTUs from metagenomes

All samples were assembled using both, MetaSPAdes v.3.15.3 and the Metaviral SPAdes option therein [43, 52]. Viruses were identified using VIBRANT v.1.2.1 [53] with the -virome option for the < 0.2 µm fraction, and VirSorter 2 [54] using -include-groups “dsDNAphage,ssDNA”. Outputs or viral scaffolds were combined and filtered to a length of >10 kb mostly representing dsDNA viruses, and CheckV v.1.0.1 [55] was run for quality checks. The viral scaffolds with attributes “medium-quality”, “high-quality”, and “complete” were kept for further analysis. Thereafter, viral scaffolds underwent dereplication at the species level in VIRIDIC v.1.0 r3.6 [56], and only one representative per species cluster was kept (preferentially a circular scaffold, otherwise the longest scaffold of a cluster). Some viral scaffolds were excluded as explained in the Supplement. Mean depth and breadth of the resulting viral operational taxonomic units (vOTUs) coverage were calculated after mapping reads to vOTUs using Bowtie2 v.2.4.5 with settings --mp 1,1 --np 1 --rdg 0,1 --rfg 0,1 --score-min L,0,-0.1 [57] for mapping of ≥90 % identical reads and following conventions of [58]. A coverage of ≥75 % (read breadth) of the viral genome was achieved by running the calcopo.rb script [59]. Coverages were analyzed using the 04_01calc_coverage_v3.rb script [45] and subsequently normalized to read depths. Moreover, vOTUs were analyzed using PhaBOX tools [60]to detect the proportion of phages and their lifestyle. All vOTUs were clustered to viral clusters (VC) at genus level using vConTACT2 v.0.9.19[61] together with a recent viral reference database (2 July 2022: https://github.com/RyanCook94/inphared/tree/b614bb92f31d55bfb0ab6180604e59838e492875) from INPHARED [62], and results were compiled using graphanalyzer v.1.5.1 [63]. PhaGCN v2.0 was used to assign viral families to vOTUs [64, 65]. Auxiliary metabolic genes (AMGs) that correspond to Class I AMGs defined by [66] were identified using AnnoVIBRANT (https://github.com/AnantharamanLab/annoVIBRANT). After gene calling using Prodigal [67], the vOTUs were further annotated using DRAM-v v.1.4.6 [68] to find AMGs related to cryosurvival. To explore virus-host interactions, vOTUs were matched to genomes from the “Sept_21_pub” database of the Integrated Phage Host Prediction (iPHoP) tool v.1.2.0 [69]. The database was complemented with dereplicated MAGs and genomes of isolated strains (see below) from this study. From the iPHoP predictions run with a confidence score cutoff of 90, the host genus with the highest confidence score was considered for visualization within Cytoscape v.3.9.0 [70]. The presence of vOTUs at five different stations (including the melt pond) based on read breadth, was correlated with the average coverage of the vOTU for all SML and SSW samples, respectively. The underlying assumption was that the higher the coverage of vOTUs at the air-sea interface, *i.e.* the more abundant they will be in the SML, the more they would be spread across the Arctic and occur at multiple stations since they likely get more easily aerosolized from the SML.

### Bacterial isolation and genome analysis

For bacterial isolation, 900 µl of seawater was added to 600 µl 50 % glycerol, inverted for mixing and stored at −80 °C. In the home laboratory, the sample water-glycerol mix was spread onto Zobell agar plates (1 g yeast extract (BD), 5 g bacto-peptone (BD), 15 g bacto agar (BD), 800 mL specific water see below, 200 mL Milli-Q water). To account for the fact that bacteria came from different aquatic sources or are potentially used to strong salinity gradients in the SML due to melting sea ice, agar plates with different water sources of different salinities were used. This was achieved by either adding Milli-Q (MQ) water, Baltic Sea (Bal) water collected from the Linnaeus Microbial Observatory [71], or Arctic Ocean (Arc) water to the Zobell plates. Incubations were performed at room temperature and 4 °C. Colonies of different morphologies were picked and clean streaked thrice. Finally, all isolated bacteria were grown at room temperature in liquid Zobell_Arc or Bal or MQ_. Stocks from all bacterial strains with 600 μl 50 % glycerol (Sigma) and 900 μl liquid overnight grown culture were stored at −80 °C. DNA from overnight cultures of bacterial isolates was extracted using E.Z.N.A Tissue DNA Kit (Omega Bio-tek, Norcross, GA, USA), eluted in 50 µl, quantified on the Qubit^®^ 2.0 Fluorometer and stored at −80 °C. In total, 13 bacterial isolates (Table S3) were sent for whole-genome Illumina sequencing to Scilifelab (Solna, Sweden) using the same platform and flowcell as mentioned above. Reads were processed as mentioned above and assembled using SPAdes v.3.15.3 with option --isolate. Prophages in MAGs and isolate genomes were searched using VIBRANT v.1.2.1. CRISPR spacer in isolate genomes were searched and extracted with the CRISPRcasFinder online tool [72] and matched to vOTUs from the metagenomes and viromes using a blastn-short algorithm with a 80 % similarity threshold.

### Prophage induction experiments and transmission electron microscopy

A prophage induction assay was performed for the bacterial strains *Psychrobacter* sp. Arc29, *Leeuwenhoekiella aequorea* Arc30, *Pseudoalteromonas distincta* Arc38, and *Flavobacterium frigidarium* Arc14 (the latter as control for a strain lacking prophages, Table S4). Due to the contamination issue when sampling SML and SSW from the gangway (see Supplement), we chose only these bacteria since they are known from the literature to be psychrophilic and/or marine bacteria. In a 48-Well plate, mitomycin C (Roche Diagnostics) as prophage-inducing reagent [73] was added in triplicates in different final concentrations: 1 µg ml^−1^, 0.5 µg ml^−1^, 0.1 µg ml^−1^, and no mitomycin C to 500 µl of overnight grown bacterial culture. To monitor bacterial growth, the optical density at a wavelength of 600 nm (OD_600_) on a FLUOstar® Omega Microplate Reader (BMG Labtech, Ortenberg, Germany) was measured once per hour. A significant OD drop under mitomycin C treatment compared to the negative control indicated prophage induction. At the end of the experiment, the culture liquid from the wells of induced prophage was collected, pooled, and subsequently filtered through a 0.2 µm syringe filter. The flow-through was further concentrated from 10 ml volume in an Amicon^®^ Ultra-4 Centrifugal Filter 50 kDa unit (Merck Millipore) and stored at 4 °C. DNA was isolated from 1 ml concentrated supernatant using Wizard PCR DNA Purification Resin and Minicolumns (both Promega, Madison, WI, USA) as previously described [74]. Since VIBRANT found two prophages in the genome of *L. aequorea* Arc30, whole-genome sequencing of the DNA from independent prophage induction experiments was performed to determine the prophage identity in the supernatant after filtration on 0.2 µm pore-size syringe filter. From one of these experiments, half of the virus supernatant after the concentration step was digested with amplification grade DNAse I (Invitrogen/Thermo Fisher Scientific, Waltham, MA, USA), for 10 minutes at 37 °C before DNA extraction to reduce host DNA contamination. Sequencing was done on a NOVASeq6000 platform by using the INVIEW Resequencing service of Eurofins Genomics (Ebersberg, Germany). After read trimming and QC as mentioned above, the phage genomes were assembled using MEGAHIT v.1.2.9 [75], and scaffolds with viral genes identified using CheckV. To be able to assemble one of the induced phages’ genomes, the trimmed reads had to be randomly reduced to 1 % using seqtk v.1.4 (https://github.com/lh3/seqtk). A circular proteomic tree was built for the induced phages using ViPTree v.4.0 [76] including genomes of related phages suggested by the tool. Intergenomic similarities with related phages were further explored in VIRIDIC [56]and protein clustering in the VirClust webtool [77]. Gene prediction and annotations were performed as explained above for vOTUs. The genome of the phage Arctus_1 was circularized in Artemis [78] and visualized using Proksee [79]. The sliding window application (window size 10000, step size 100) within Proksee was used to look at the GC content distribution of the genome. To check for the biogeography of the induced prophages, their genomes were blasted against the IMG/VR viral nucleotide database v.4 [80] with e-value of 1e-5.

Phage supernatants were examined by transmission electron microscopy (TEM) using negative staining as in [57]. Phages were loaded on pre-discharged 200 mesh copper grids covered with carbon film (Agar Scientific Ltd., Stansted, UK), and stained with 2 % w/v uranyl acetate. Images were taken with the FEI Tecnai 12 G2 BioTWIN microscope. Capsid diameter and tail length were measured from TEM images using ImageJ v.1.53t according to previously published protocol [81]. A successful cross-pole infection of Arctic *Pseudomonas* sp. G11 with lysogenic *Pseudomonas* phage vB_PaeM-G11 isolated from an Antarctic strain was recently shown [82], and inspired us for a similar cross-infection experiment. The induced Arctic phages were tested on an *L. aequorea* strain CCUG 50091^T^ [83] isolated from Antarctic seawater and ordered from the Culture Collection of University of Gothenburg, Sweden. However, no evidence for cross-infection could be found for this phage-host pair in plaque assay or liquid culture.

### Statistics

Unpaired, non-parametric Mann-Whitney U test for comparing means of the Shannon-Wiener index was carried out in Graphpad Prism v.10. Before the correlation analysis of vOTU coverage with distribution at stations, outliers were removed using the Robust regression and Outlier removal (ROUT) method with Q = 1, as recommended in [84] and implemented in Graphpad Prism v.10, where also linear regressions where drawn. Spearman *r* correlation coefficient was applied, and groups for the different stations were compared using Kruskal-Wallis test with Dunn’s multiple comparison test.

## Results

### Taxonomic profiling of prokaryotes and metabolic insights from Arctic metagenome-assembled genomes

According to the taxonomic profiling with the tool mOTUs, the melt pond was dominated by *Bacteroidota*, and particularly by a *Flavobacterium* sp. [genome reference from mOTUs ext_mOTU_v3_22321], which was phylogenetically related to the melt pond-derived *Nonlabens* sp. (MAG_04 from our study) (Figure 2a, Figure S1) and had a relative abundance of 60.1 % and 35.1 % in the 0.2 – 5 µm fraction of the SML and SSW, respectively. A relative abundance of 21.6 % and 32.9 % from SML and SSW of the same size fraction remained unassigned. Other OTUs with >1 % relative abundances in the melt pond SML and SSW samples from the 0.2 – 5 µm and >5 µm fraction were *Oceanospirillaceae* species, *Cellvibrionales* species, *Betaproteobacteria* bacterium MOLA814, *Rickettsia* sp., *Octadecabacter arcticus*, *Pelagibacteraceae* species, and *Polaribacter* sp. (Table S5). Open water samples were overall more diverse, as the Shannon-Wiener index indicated, with a range of 1.1 – 2.3 (*n* = 4) for the melt pond, and a range for open water samples of 2.4 – 3.7 (*n* = 8). The difference between the two means was significant (two-tailed Mann-Whitney U-test, U value = 0, *p* = 0.0040, Figure 2c). However, a clear limitation is that the four melt pond samples were just different filtered fractions and all from the same melt pond, thus only providing a first glance at the low diversity of bacteria in melt ponds. Arctic open water samples contained less *Bacteroidota* (max: 16 %) and more *Pseudomonadota* (max: 37.1%) compared to the melt pond (*Bacteroidota* max: 60.1 %, *Pseudomonadota* max: 7.9 %), but a similar number of unassigned taxa. While the melt pond was mainly dominated by a single *Flavobacterium* sp. up to 60.1%, open water samples contained up to 19.6 % relative abundance of up to 45 different species of *Flavobacteriaceae*.

**Figure 2:**
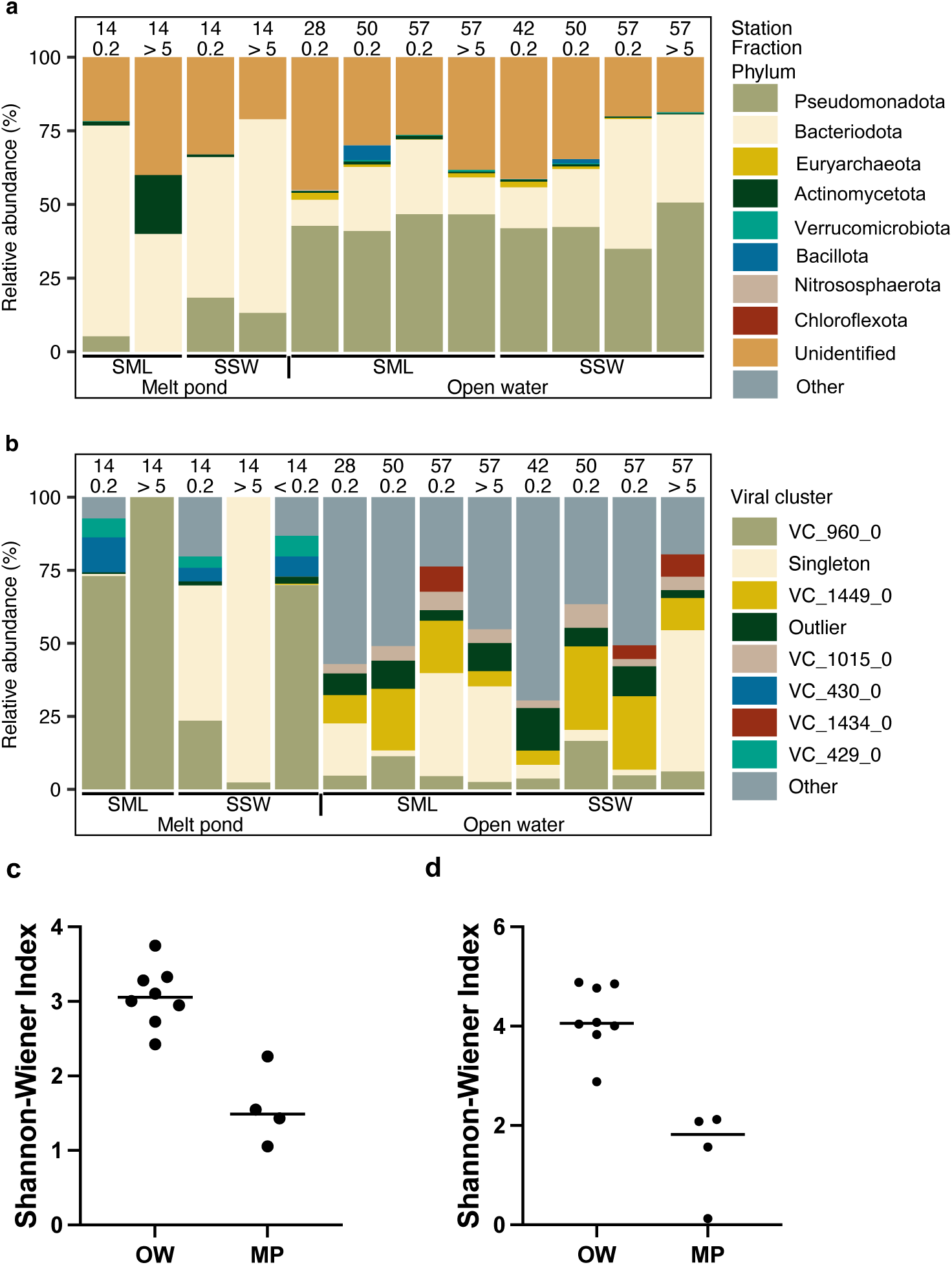
Relative abundances of prokaryotes and viral clusters (VCs) in surface microlayer (SML) and subsurface water (SSW) stations sampled across the Central Arctic. Community structure based on mOTUs depicting the nine most abundant prokaryotic phyla while grouping the remaining phyla as “Other” (a). Community structure based on VCs depicting the nine most abundant viral groups while grouping the remaining clusters as “Other” (b). Fraction 0.2 refers to the filtered fraction between 0.2 µm to 5 µm pores sized filter membranes, whereas fraction 5 is the ≥ 5 µm pore size filtered sample. Alpha diversity of the mOTUs for melt pond (MP) and open water (OW) samples estimated according to the Shannon-Wiener index for mOTUs (c) and vOTU communities (d). The line indicates the median.

A total of 182 metagenome-assembled genomes (MAGs) were binned and dereplicated to 68 MAGs (Table S6). After further exclusion of likely contaminants, resulting in 50 MAGs, three MAGs were assigned to archaea (phyla *Thermoproteota* and *Thermoplasmatota*). Others were assigned to the bacterial phyla *Actinomycetota* (5), *Bacteroidota* (19), *Proteobacteria* (18), SAR324 (2), and *Verrucomicrobiota* (3). From 13 bacterial isolates, 8 dereplicated genomes were assigned to *Bacteroidota* (3), *Pseudomonadota* (4), and *Bacillota* (1). The average estimated completeness of these genomes was 91 ± 8 % (mean ± STD, *n* = 58, Fig. S2a) and based on the metabolic potential, the archaeal MAGs clearly diverged from the bacterial MAGs (Fig. S2c). The estimated genome size was similar for both the MAGs in the melt pond and open water and was 2 Mbp on average (SD = 0.51, *n* = 58, Fig. S2b). Predictions on the index of replication (iRep) indicated that none of the bacterial isolates and only five MAGs, i.e., *Nonlabens* sp. (MAG_04), *Polaromonas* sp. (MAG_07), a *Spirosomaceae* bacterium (MAG_08), *Aquiluna* sp. (MAG_09), and *Burkholderiaceae* bacterium (MAG_34) were actively replicating in the melt pond sampled at Station 14 (iRep range 1.4 – 2.2, Table S7). Of these, *Aquiluna* sp. had the highest coverage of reads (up to 123 x) indicative of the genome’s abundance. In open water samples, iRep values indicated replication indices for 46 MAGs, with the highest values being found for *Polaribacter* sp. (MAG_101, iRep = 5.9 and 3.4 at Station 28 & 42), *Patiriisocius* sp. (MAG_75, iRep = 5.1 – 6.0 at Station 57), and a *Flavobacteriaceae* bacterium (MAG_123, iRep = 4.0 – 4.9 at Station 57). *Candidatus* Pelagibacter (MAG_10), a Gammaproteobacteria bacterium (MAG_161), and a *Porticoccaceae* bacterium (MAG_168) were abundant on various stations but not among the most active bacteria based on iRep predictions. The bacterial strain *L. aequorea* Arc30, isolated from SML of Station 42, for which two prophages could be successfully induced (see below), had a maximum coverage of 0.2 x and thus no predicted iRep, suggesting that it was beyond detection limit in the metagenomes and very low abundant. Metabolic potential was analyzed for 29 MAGs, showing that in both the melt pond and open water heterotrophy prevailed due to the absence of the major carbon fixation pathways, and prokaryotes obtained their energy from the oxidation of organic compounds (Fig. 3). Taxonomic profiling of the prokaryotic community based on MAGs is shown in Figure S3a&b.

**Figure 3:**
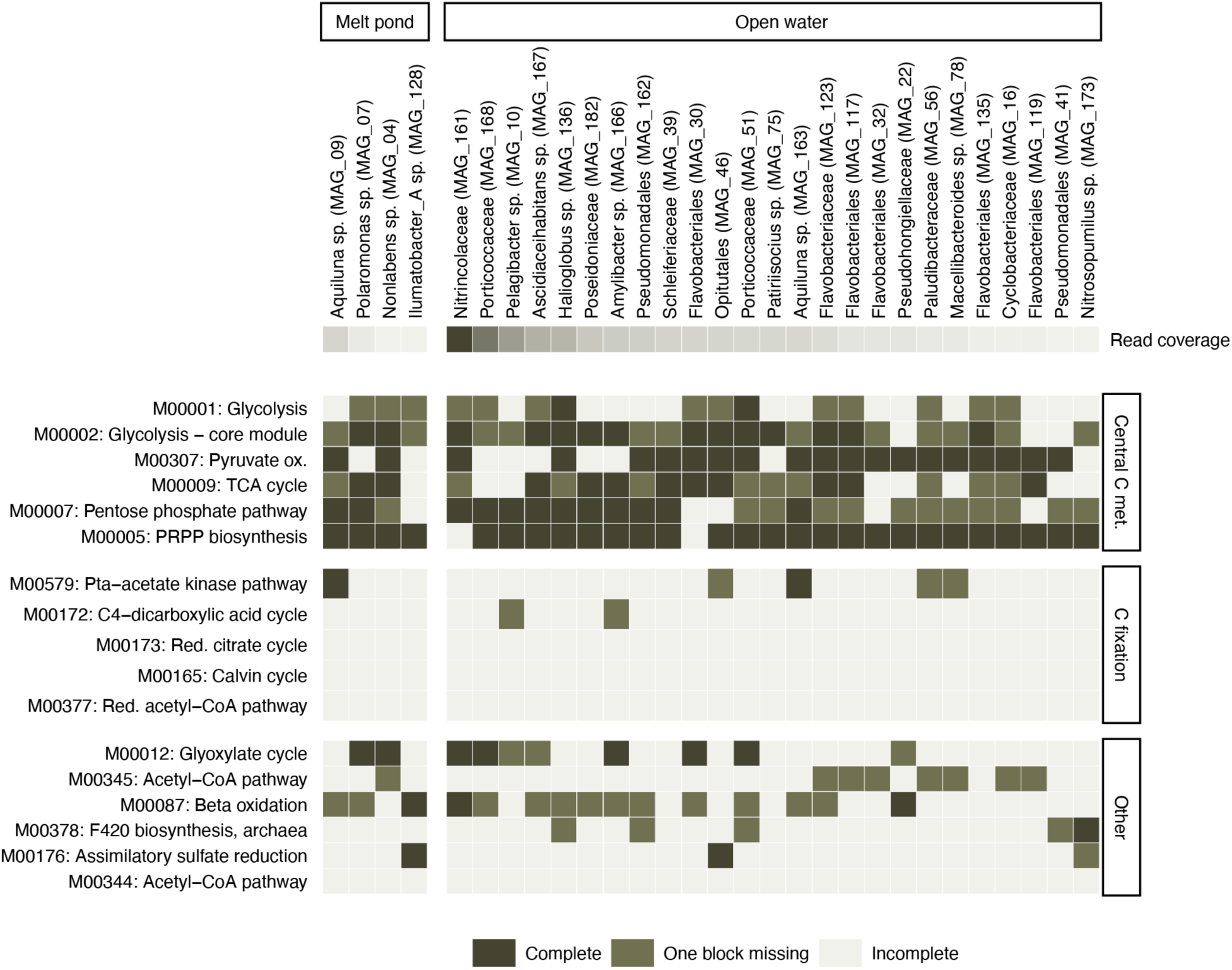
Metabolic functions deduced from the MAGs. The MAGs were ordered based on abundance (read coverage) while showing the most abundant MAGs for both the melt pond (*n* = 4) and the open water (*n* = 25). The color code depicts the completeness of the KEGG modules, divided into three categories.

### Viral community, lifestyle, auxiliary metabolic genes, and dispersal from the SML

In total, 1154 dereplicated vOTUs were detected, of which PhaMer [85] predicted 66.7 % (770) to be phages. Of those, PhaTYP [86] predicted 79.6 % and 20.4 % to be virulent and temperate phages, respectively (Table S8). The > 5 µm and 0.2 – 5 µm filtered fractions of the melt pond SML were dominated by VC 960_0 (containing vOTU1148 and vOTU1151) corresponding to a Skunavirus-like family and sharing protein similarities with marine Nonlabens phage P12024S (GenBank #<u>JQ823122</u>) [87] (Figure 2b, Figure S3c, Table S9) with 100 % and 54.1 % relative abundance, respectively. The melt pond SSW showed a prevalent vOTU identified as Singleton in vConTACT2 assigned to *Metaviridae* family and had 97.7 and 46.0 % relative abundance on the > 5 µm and 0.2 – 5 µm pore-sized filters, respectively (Figure 2b, Figure S3c). Both SML and SSW, contained the VC 429 and VC 430, which encompass vOTUs sharing protein similarities with the siphovirus Flavobacterium phage 11b (GenBank #<u>AJ842011</u>), previously isolated from Arctic sea-ice [14]. In total, only 10 different vOTUs including the two very abundant vOTUs described above reached a relative abundance > 2 % in the melt pond, which shows that only few viruses thrive there and adapt to the low diversity of hosts. In the open water, the alpha diversity was higher than in the melt pond (two-tailed Mann Whitney U-test, U value = 0, *p* = 0.0040, Figure 2d), and vOTUs not assignable to known families were most abundant and found at every station (Figure S3c), being in line with the novelty of vOTUs as inferred from clustering proteins with viruses from a RefSeq database (Figure 4, Table S9).

**Figure 4:**
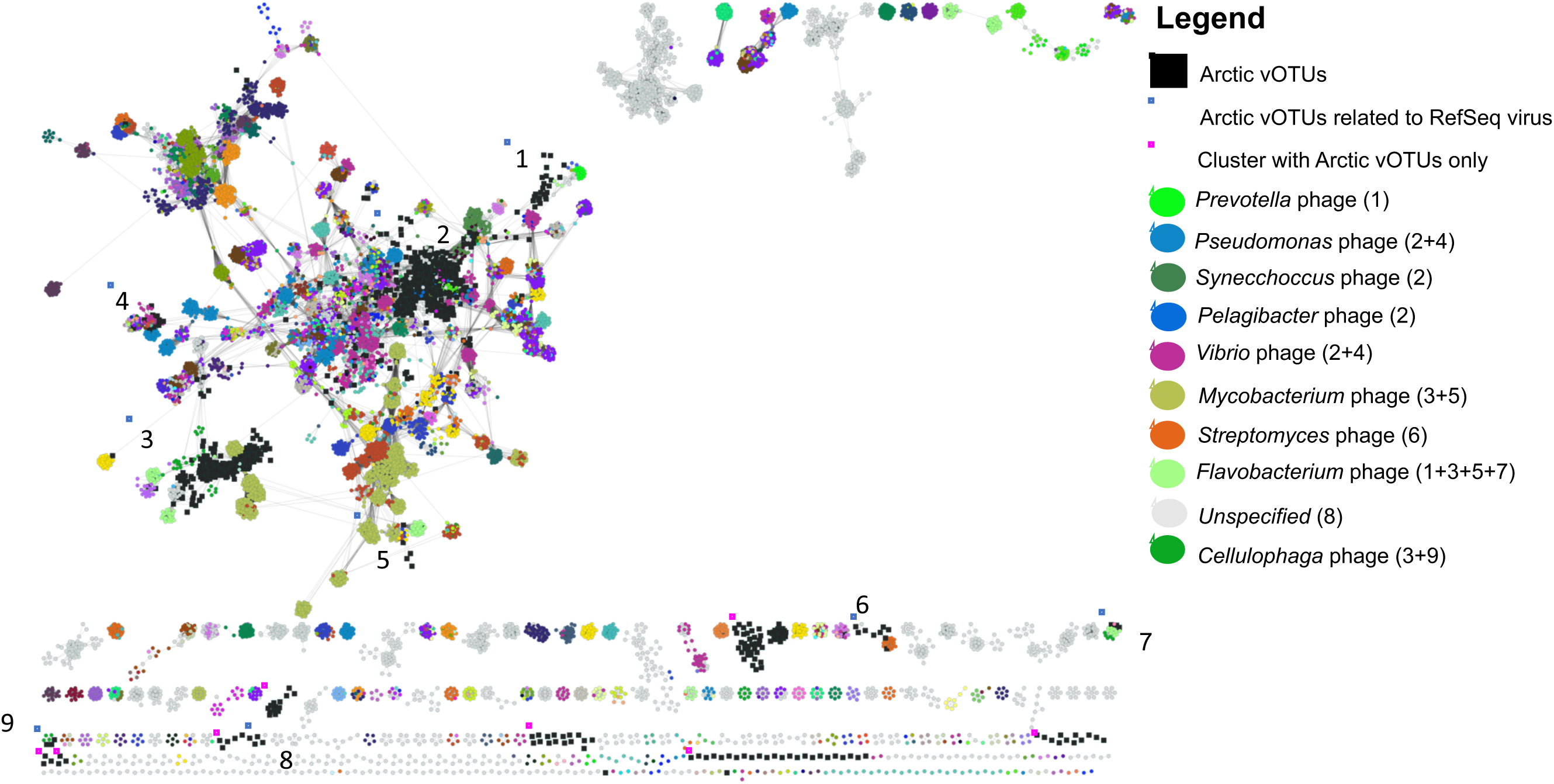
Network of viral OTUs from this study clustered with viral genomes from the RefSeq database (July 2022) based on shared proteins. Blue, numbered frames show Arctic vOTUs clustering with known phages, with the most important clustering partners including host name shown in the legend with frame number in parentheses. Pink frames indicate where Arctic vOTUs form clusters with each other.

Host prediction using iPHoP could link 98 vOTUs to a host at a minimum confidence score of 90 (Table S10, Figure 5a) and of these, five vOTUs were predicted to have an archaeal host. The most common order of bacterial hosts was *Flavobacteriales*, to which 24 vOTUs were assigned. CRISPR analysis revealed that 8 of the 13 bacterial strains had CRISPR arrays as part of the adaptive immune system of prokaryotes, but only one CRISPR spacer from *Flavobacterium* sp. Arc2 matched a vOTU (coverage in melt pond = 214 x) assembled from the melt pond virome (Table S11). This vOTU was also assigned to *Flavobacterium* as the host in iPHoP.

**Figure 5:**
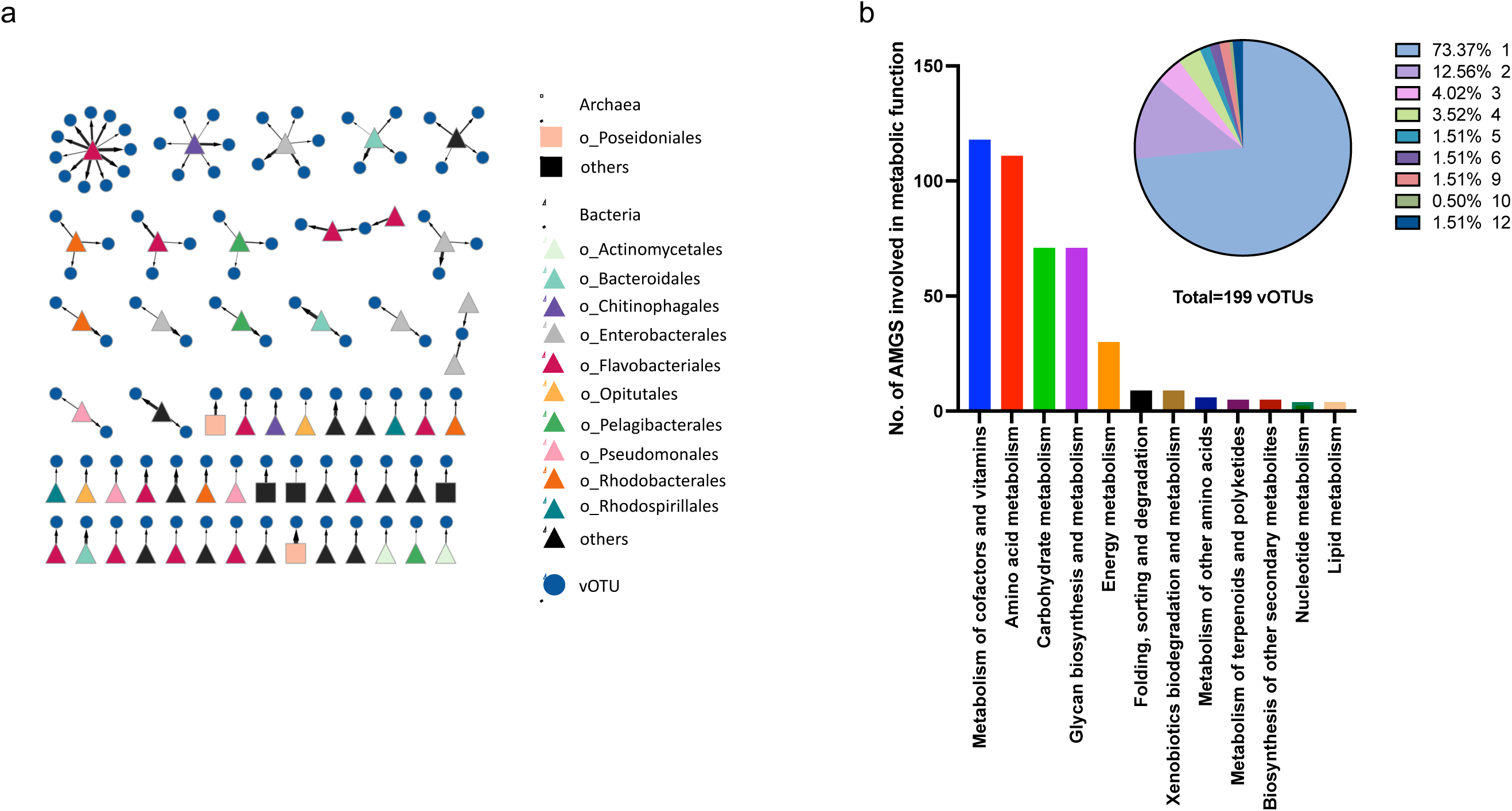
Virus-host matches and AMGs. Host assignment for 96 Arctic vOTUs derived from iPHoP prediction (a). Minimum confidence score = 90; the thicker the arrow, the higher the confidence. For further details, please see Table S10. Number of auxiliary metabolic genes involved in metabolic functions (b); 199 vOTUs carried 87 unique AMGs (354 in total) with involvements in different metabolic pathways (Table S12). The pie chart shows the percentage of vOTUs carrying one or more (up to 12) AMGs (Table S13).

Prophages were mostly found in SML isolate genomes but were rare in MAGs (3 in bacteria, 2 in archaea, Table S4). Interestingly the archaeal MAGs 172 and 175 belonging to Marine Group II archaea within *Euryarchaeota* and sampled from stations 42 and 50 both carried similar prophages with one auxiliary metabolic gene (AMGs) for glycine cleavage system H protein gcvH (KEGG database <u>K02437</u>) and glucose-1-phosphate thymidylyltransferase rfbA, rffH (KEGG database <u>EC:2.7.7.24</u>., <u>K00973</u>) involved in the biosynthesis of thymidine-linked sugars. In the remote Central Arctic, prophages might have an important function in assisting metabolic functions in these Archaea. Several prophages were found in bacterial isolate genomes from the SML, e.g. *Burkholderia vietnamiensis* Arc4 carried 8 prophages, while *Bacillus pumilus* Arc15, *Psychrobacter* sp. Arc29, and *L. aequorea* Arc30 each carried 2 prophages, and *P. distincta* Arc38 and *Alcaligenes phenolicus* Arc10 (the latter likely being a contaminant associated with gangway sampling) both carried 1 prophage. All prophage annotations are reported in Table S4.

Using AnnoVIBRANT, we identified that 199 vOTUs (17.2 % of all vOTUs) carried 87 unique AMGs (354 in total) with involvements in various, sometimes multiple metabolic pathways (Figure 5b, Table S12). Of the AMG-carrying vOTUs, 26.6 % carried more than one AMG and up to twelve in total (Figure 5b, Table S13). Most abundant AMGs had predicted functions in amino acid metabolism (111), mainly in cysteine and methionine metabolism, in metabolism of cofactors and vitamins (118), mainly for porphyrin, as well as in glycan polymer and carbohydrate metabolism (both 71). AMGs related to cryoprotection of the host were also identified. Sixteen vOTUs carried the AMG glycerol-3-phosphate cytidylyltransferase or *tagD* [Enzyme Commission number, <u>EC:2.7.7.39</u>]. This gene encodes for an enzyme involved in the production of teichoic acids [88], which in turn benefit the freeze tolerance in bacteria [89, 90]. Three further AMGs were annotated as ice-binding-like [Pfam/InterPro database, <u>PF11999.11],</u> which as gene family includes ice-binding proteins (IBPs). In addition, vOTU encoded genes related to cell wall metabolism were found, namely 199 AMGs for N-acetylmuramoyl-L-alanine amidase [<u>EC:3.5.1.28</u>], 140 AMGs for zinc D-Ala-D-Ala carboxypeptidase [<u>EC:3.4.17.14</u>], and 35 AMGs for peptidoglycan LD-endopeptidase CwlK [<u>EC:3.4.-.-,]</u>. All DRAMv annotations of vOTUs are shown in Table S14.

By assuming that vOTU distribution through the Arctic would be mediated by the depth of the layer where the vOTU was found in (the closer to the atmosphere, the more prone to aerosolization, Figure 6b), linear regressions were performed for vOTU abundance based on read coverage and the number of stations a vOTU was present based on read mapping. We found positive correlations for both vOTUs from SML and SSW but a steeper slope for the SML (1.254) compared to the SSW (0.4673, Figure 6a) indicating that vOTU abundance changes in the SML are more relevant for spread to different stations than changes in the SSW. The goodness of fit was better for SML (R^2^ = 0.49) than for SSW (R^2^ = 0.25) data. Spearman r for the positive correlation between vOTU abundance and number of stations where vOTUs were detected was 0.78 (*n* = 375) and 0.38 (*n* = 649) for SML and SSW, respectively. Statistics for group comparisons are shown in the supplement material (Figure S4).

**Figure 6:**
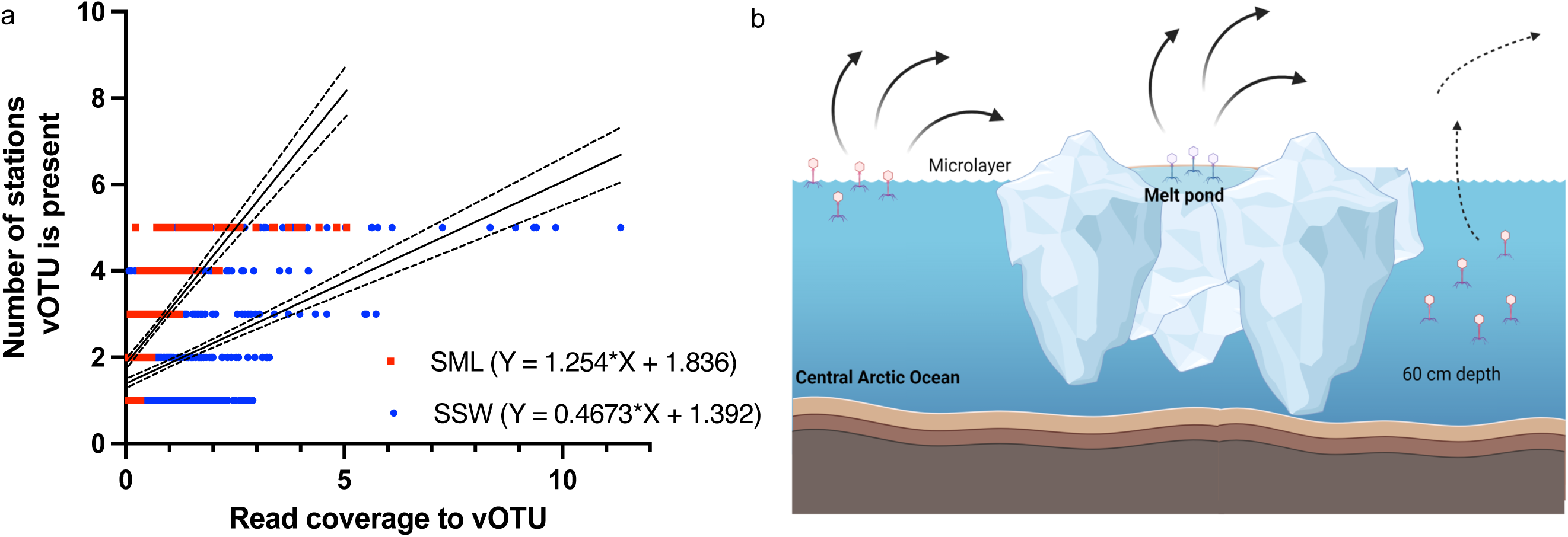
Correlation of vOTUs coverage with presence at stations. Linear regressions (mean with confidence intervals) showing the positive correlation between the vOTU average read coverage for SML and SSW with the number of stations a vOTU was present based on read breadth (a). The concept figure illustrates the intensified spread of viruses from surface films of Arctic Ocean and melt pond air-sea interface compared to deeper surface water irrespective of the higher abundance of phages in the SSW (b).

### Prophage induction from strain *L. aequorea* Arc30

The *L. aequorea* Arc30 culture showed a drop in OD_600_ ~4 hours after addition of mitomycin C (1 and 0.5 µg ml^−1^) but not in the 0.1 µg ml^−1^ and no mitomycin C condition, and together with phage appearance in TEM, indicated prophage induction to the lytic cycle (Figure 7a). This OD drop was reproducible in several independent induction experiments. Within the genome of *L. aequorea* Arc30, VIBRANT predicted two prophages of 36.4 kb and 44.1 kb. Mapping of reads from the induction’s supernatant to both prophage regions of the host, showed that both prophage regions had a very high coverage, indicating induction and thus activity of both prophages (Figure S5). Whole-genome sequencing of the phages from the supernatant after induction revealed that the 44.1 kb genome was circular (Figure 7b), had a length of 42.7 kb, 34.91 % GC content, and 75 open reading frames. Genes encoding for structural proteins were found in above average GC regions using a sliding window, while DNA replication/recombination/repair and packaging genes were found in below average GC regions (Figure 7b). The integration site of the 44.1 kb prophage was a host tRNA for methionine (codon CAT), the phage itself carries a tRNA for tryptophan (codon CCA). For the induced 44.1 kb prophage (42.7 kb when excised) we propose the name Leeuwenhoekiella phage vB_LaeS_Arctus_1. Arctus_1 is distantly related to known *Polaribacter* phages based on shared proteins (Figure 7d, Figure S6) but the intergenomic similarity to these phages is < 1.3 % (Figure S7). We propose “Leeuwenvirus” and “Leeuwenvirus arctus” as new genus and species name for Arctus_1, respectively. As a viral family name, we propose *Neustonviridae* because the virus was isolated from the SML, where the neuston organisms reside.

**Figure 7:**
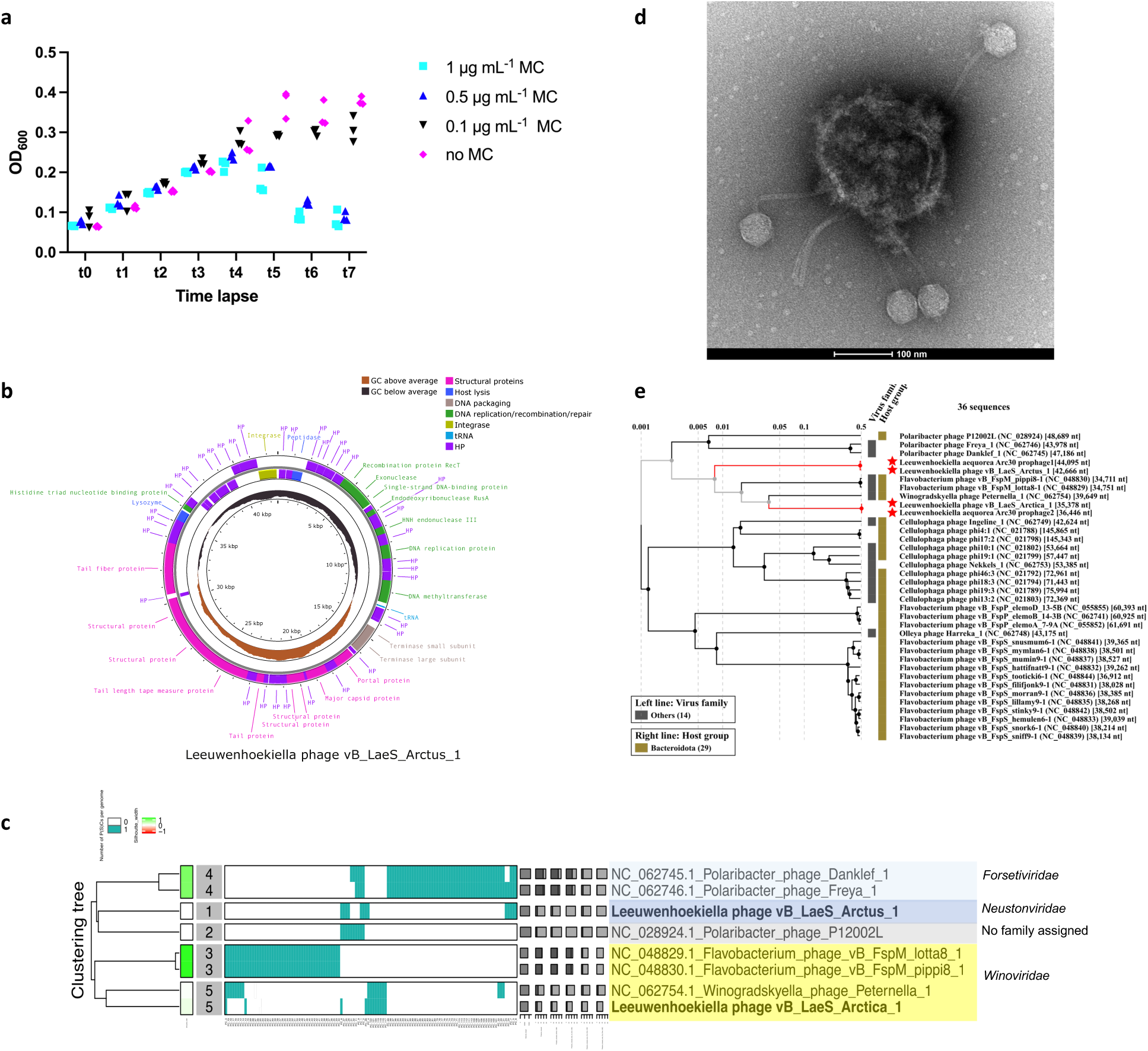
Induction of two *Leeuwenhoekiella* prophages. Growth curve of the strain *Leeuwenhoekiella aequorea* Arc30 indicating prophage induction by mitomycin C treatment at t4 (a). The architecture of the 42.7 kb circular genome belonging to the induced Leeuwenhoekiella phage vB_LaeS_Arctus_1 (b). Protein clustering tree of Arctus_1 and Arctica_1 with related phages (c). TEM imaging at 68k magnification revealed siphoviruses attached to a lysing host cell in the supernatant of the bacterium *L. aequorea* Arc30 after treatment with mitomycin C (d). A proteomic tree showing the placement of Leeuwenhoekiella phage vB_LaeS_Arctus_1 (e).

The genome of the other, 36.4 kb prophage (35.4 kb when excised) was found in three independent assemblies, but the genome could not be circularized and might not be complete. The linear phage shares nine core proteins with Winogradskyella phage Peternella_1 (GenBank #<u>NC_062754</u>) isolated from surface water of the North Sea [91] (Figure S6), hence we suggest to assign it to the same family, *Winoviridae*. The phage genome has 42.0 % GC content and 43 open reading frames. We propose the name Leeuwenhoekiella phage vB_LaeS_Arctica_1 and “Leeuwenhoekiellavirus” and “Leeuwenhoekiellavirus arctica” as new genus and species name for Arctica_1. Arctus_1 and Arctica_1 share 4.2 % intergenomic similarities with each other (Figure S7). Shared core proteins and functional annotations for the phages are shown in Table S15.

TEM imaging revealed siphoviruses (head and non-contractile tail) in the supernatant of induced *L. aequorea* Arc30 lysogen, with the phages attached to the lysed host cell (Figure 7c). The phages had a mean ± standard deviation diameter of the capsid of 50.4 ± 3.0 nm (*n* = 24) and a tail length of 157.7 ± 12.2 nm (*n* = 16, Table S16). Arctus_1 had a tape measure protein of 978 amino acids length (for sequence see Supplement material). According to a formula proposed earlier [92], the calculated tail length of that phage would be 154.8 nm, which is very close to the length of the phage observed in TEM. Arctica_1 had no identifiable tape tail measure protein. The genomes of Arctus_1 and Arctica_1 had hits against genomes from uncultivated viruses assigned to the class *Caudoviricetes* in the IMG/VR database originating from marine, freshwater, saltwater ecosystems from around the globe, including Arctic and Antarctic ecosystem, and additionally matches against genomes of other *Leeuwenhoekiella* sp. strains (Table S17).

## Discussion

The aim of this study was to investigate viral-bacterial interactions in atmosphere-close aquatic ecosystems of the Central Arctic. Our study shows that viral and bacterial diversity was lower in the melt pond compared to the oceanic samples. Melt ponds form due to melting sea ice and snow in Arctic summer and are typically not connected with the ocean underneath the ice. Due to their shallow depth, melt pond water is prone to more freeze-thaw cycles, with the lowest temperature at the surface of the pond (where the SML exists) [93] adding an extra burden to therein living organisms and viruses and probably restricting their diversity. A lower alpha diversity as well as a different community composition of microbial eukaryotes in melt ponds compared to seawater was previously reported [94]. We noticed that only two melt pond viruses became very abundant in filtered fractions > 0.2 µm suggesting that they responded to and thrived well on the few available hosts. Melt ponds selected for *Flavobacteria* (*Bacteroidota*), *Betaproteobacteria*, and *Alphaproteobacteria* depending on factors such as if melt ponds are open or closed as well as the salinity of the water [95, 96]. The melt pond studied here was open when sampled and primarily enriched in *Pseudomonadota* such as *Polaromonas* sp. (MAG_01), *Bacteroidota* such as *Flavobacterium* sp. (Arc3) and *Nonlabens* sp. (MAG_04), as well as *Actinomycetota* such as *Aquiluna* sp. (MAG_03). Due to their disconnection from the ocean, the major source for biological input, including viruses, into melt ponds probably is melting ice and atmospheric exchange. Sea ice algae, such as the diatom *Melosira arctica,* which was observed during this cruise, can show large aggregates in melt ponds (reviewed by [97]). The SML from the sampled melt pond was highly viscous (personal observation), suggesting that algae-derived extracellular polymeric substances (EPS) were likely present. This would also explain the predominance of the family *Metaviridae*, which includes retrotransposons and reverse-transcribing viruses targeting eukaryotes [98] and thus potentially algae. However, while we provide first insights into viruses from melt ponds, a clear limitation of our study is that only a single melt pond could be sampled, leading to the question, if different melt ponds contain similar or different viral-bacterial communities, which must be addressed in future investigations.

Virus-encoded AMGs related to cryosurvival have previously been described for viruses from the Southern Ocean [99], including genes related to cell wall polymer and extracellular polymeric substance (EPS) production, or cold shock genes, the latter being also detected in viruses from Arctic glacial ice [100]. IBPs have been described in many bacteria [101, 102], where they prevent damage from ice crystals by binding to ice surfaces and inhibiting their growth. An antifreeze protein belonging to a subset of the IBPs has been found in viruses derived from metagenomes of the Southern Ocean [99]. In addition to IBPs, we also found AMGs encoding for glycerol-3-phosphate cytidylyltransferase, which has a function in the biosynthesis of teichoic acid, a cell wall component of Gram-positive bacteria. Constituents of teichoic acids undergo chemical interactions that allow liquid water to be maintained within the cell wall and the immediate extracellular space [89, 90] thereby providing cryoprotection. Teichoic acids can be secreted into extracellular spaces, for instance within biofilms [103]. Possessing these AMGs might increase the chances of host viability and thus benefit viral replication and dissemination under harsh conditions. Having antifreeze and freeze-tolerance genes in the SML can represent a meaningful advantage, namely that the bacterial hosts acquire cryoprotection in habitats exposed to freezing. When sea water freezes, the formation of ice typically initiates at the surface due to direct exposure to cold air, gradually expanding downward as the freezing process continues.

We noticed that vOTU distribution to many stations was more strongly positively correlated with average coverage for SML vOTUs than for SSW vOTUs, despite the average coverage of an vOTU overall being higher in SSW. An explanation could be that viral accumulation in SML facilitates their aerosolization and atmospheric distribution so that viruses can spread more easily through the Arctic. It follows that aerosolization might be easier for low to medium abundant vOTUs in the SML than for higher abundant ones in the SSW. Whether such a correlation exists for other oceans, however, remains to be understood. Aerosolization of viruses from SML is a known feature [15, 16, 104] and is heavily mediated by bubbles that rise to the surface and scavenge viruses [105, 106]. Recent work with tank experiments has shown that virus transfer leading to aerosolization from water surfaces is related to the size of bubbles (the smaller, the more viruses) and their originating depth (the deeper, the more viruses are scavenged on the way to the surface) [107].

We found two inducible, i.e. active prophages for the low-abundant Arctic SML strain *L. aequorea* Arc30. In contrast, results from Baltic Sea SML showed, that an inducible phage occurred in one of the most abundant bacterial strains, *Alishewanella* sp. being additionally hunted by lytic phages [108]. Flavobacteria were previously found in the SML from different oceans [109, 110], represent frequent hosts for lytic and temperate phages in the SML [108], and were important predicted viral hosts in the Arctic in this study. Lysogeny is often prevalent or dominant in polar environments [82, 111, 112]. A switch from lysogenic to lytic viral lifestyle mainly happens when bacterial production increases as shown for samples from the Southern Ocean [111]. However, even in environments of low productivity, viruses can be active and infect key prokaryotes as shown by a recent study on virioplankton from under the Antarctic Ross Ice Shelf [113]. Similarly, in a previous investigation on virioneuston activity from the Arctic and Antarctica, the lytic viral strategy dominated over lysogeny as deduced from calculations on flow cytometry counts of virus-like particles and prokaryotes after mitomycin C treatment of water samples [22], matching our observation of (inducible) prophages in the Central Arctic’s SML. How often prophage inductions happen spontaneously in nature is another question that warrants further investigation. The SML compared to the pelagic ocean may be a more virally active ecosystem in the Arctic, particularly in summer, when long hours of solar and UV radiation hit the air-sea boundary and naturally enhance phage induction [114]. This might be counter-balanced by salinity effects as high but not low salinity was associated with higher titers of the marine phage varphiHSIC infecting *Listonella pelagia,* and elevated salinity could influence the switch from lysogenic to lytic cycle [115]. Melting sea ice results in a less saline SML, therefore potentially favoring lysogeny. As the samples were retrieved close to ice edges (Fig. 1b), this could explain for the many SML-associated (compared to SSW) prophages detected in this study. The SML’s salinity is prone to fluctuations [116], immediately responds to freshwater fluxes as known from influences of rainwater [117], while at other times, the SML is more saline than the underlying water due to evaporation effects [118]. Like in any other ocean [119], the SML in polar regions is usually colder than the SSW, which is known as the cool skin layer effect [120]. We thus speculate that the lifestyle of the Arctic virioneuston in contrast to the virioplankton is under a stronger pressure to adapt to the cold temperature and freezing/melting ice conditions in the environment with limited host diversity.

## Conclusions

In conclusion, our data shed light on virus-host interactions in aquatic ecosystems of the Central Arctic Ocean, specifically in the aquatic environment interfacing with the atmosphere. The melt pond viral community differed from the oceanic viruses and responded to the availability of few hosts. Flavobacteria play a crucial role as viral hosts in both aquatic environments; however, their diversity is more pronounced in the ocean. All prophages (except for one) were found in MAGs or isolate genomes derived from the SML, and for the bacterial strain *L. aequorea* Arc30 we could prove the prophage’s activity. Lysogeny might prevail in the SML at times when influences of melting ice make the SML less saline. From the SML, vOTUs get more easily distributed across the Arctic, and some vOTUs from the first centimeters of the water surface potentially contribute to freezing tolerance of their hosts by carrying relevant AMGs. We conclude that in the Central Arctic, viruses have developed sophisticated strategies for adapting to challenging conditions. Despite the anticipated increase in ice loss and significant perturbations in the coming decades, it appears that viruses are well-equipped to persist and thrive in this ecosystem.

## List of abbreviations

AMG: Auxiliary metabolic gene
CRISPR: Clustered Regularly Interspaced Short Palindromic Repeat
EPS: Extracellular polymeric substances
IBP: Ice-binding protein
iRep: Index of replication
KEGG: Kyoto Encyclopedia of Genes and Genomes
MAG: Metagenome-assembled genomes
SML: Sea-surface microlayer
SSW: Subsurface water
TEM: Transmission electron microscopy
vOTU: Viral operational taxonomic unit

## Supplementary Information

Additional file 1: A .docx file with seven additional figures and more text information

Additional file 2: A .xlsx file including 18 tables. The titles of the tables are TableS1_Sampling_info (Table S1), TableS2_MAG_breadth (Table S2), TableS3_Bacterial_isolates (Table S3), TableS4_Prophages (Table S4), TableS5_results_motus (Table S5), TableS6_MAGs (Table S6), TableS7_iRep (Table S7), TableS8_PhaMer_Phatyp (Table S8), TableS9_vOTU_VCinfo (Table S9), TableS10_Host_prediction (Table S10), TableS11_CRISPRmatches (Table S11), TableS12_AMGnumbers (Table S12), TableS13_AMGonvOTU (Table S13), TableS14_vOTU_annotations (Table S14), TableS15_Phage_Annotations (Table S15), TableS16_TEM measure (Table S16), TableS17_IMGVR hits (Table S17), TableS18_Bioproject_PRJNA950101 (Table S18).

## Declarations

### Ethics approval and consent to participate

Not applicable for the present study.

### Consent for publication

Maria Chechik, University of York, provided consent to publish her TEM images.

### Availability of data and material

The sequencing datasets supporting the conclusions of this article are available at NCBI’s Sequence Read Archive under Bioproject ID PRJNA95010, Biosample accessions SAMN33983821 – SAMN33973845. Bacterial isolate genomes are stored at Genbank under Biosample accessions SAMN35056136 – SAMN35056148. The 182 MAGs are stored under Biosample accessions SAMN34587563 – SAMN34587744.

The vOTUs is stored under Biosample SAMN33983825. The genomes of Arctus_1 and Arctica_1 can be found at Genbank XXX (status: submitted but not released). For further information (e.g. run accessions) please see Table S18. TEM images are available at Figshare, doi: 10.6084/m9.figshare.25343113.

### Authors’ contributions

JR conducted sampling, experimental work, data analysis, and wrote the first draft of the manuscript, and together with KH isolated bacteria, contributed to prophage analysis and provided funding for this work. GW conducted metabolic analysis on MAGs and provided community plots. JW contributed to prophage induction experiments. AA provided TEM images of the induced phage. All authors contributed to editing and writing of the manuscript.

## Acknowledgements

The Swedish Polar Research Secretariat (SPRS; https://polar.se) organized and supported the SAS-Oden 2021 expedition with IB Oden in the Central Arctic Ocean. This expedition was the Swedish contribution to the International “Synoptic Arctic Survey” (SAS; https://synopticarcticsurvey.w.uib.no/). We thank the Master and crew of IB Oden for expertly undertaking the SAS-Oden 2021 expedition. We particularly thank the chief scientist Pauline Snoeijs Leijonmalm and Maria Samuelsson for enabling and supporting the project. We further acknowledge help by the science party, and in particularly Claudia Morys, Emma Svahn, Julia Muchowski, Sonja Murto, and John Prytherch for assistance during SML sampling. We thank Maria Chechik for help with TEM imaging. We thank Leonie Jaeger for discussions on the cool-skin layer effect. Data handling was enabled by resources provided by the National Academic Infrastructure for Supercomputing in Sweden (NAISS) and the Swedish National Infrastructure for Computing (SNIC2020-16-49) at UPPMAX partially funded by the Swedish Research Council through grant agreement no. 2022-06725 and no. 2018-05973. We like to thank Pavlin Mitev at UPPMAX for support. We further acknowledge support from the National Genomics Infrastructure (NGI) in Genomics Production Stockholm funded by Science for Life Laboratory, the Knut and Alice Wallenberg Foundation, and the Swedish Research Council.

