## Supplementary Material for "Viral challenges and adaptations between Central Arctic Ocean and atmosphere"

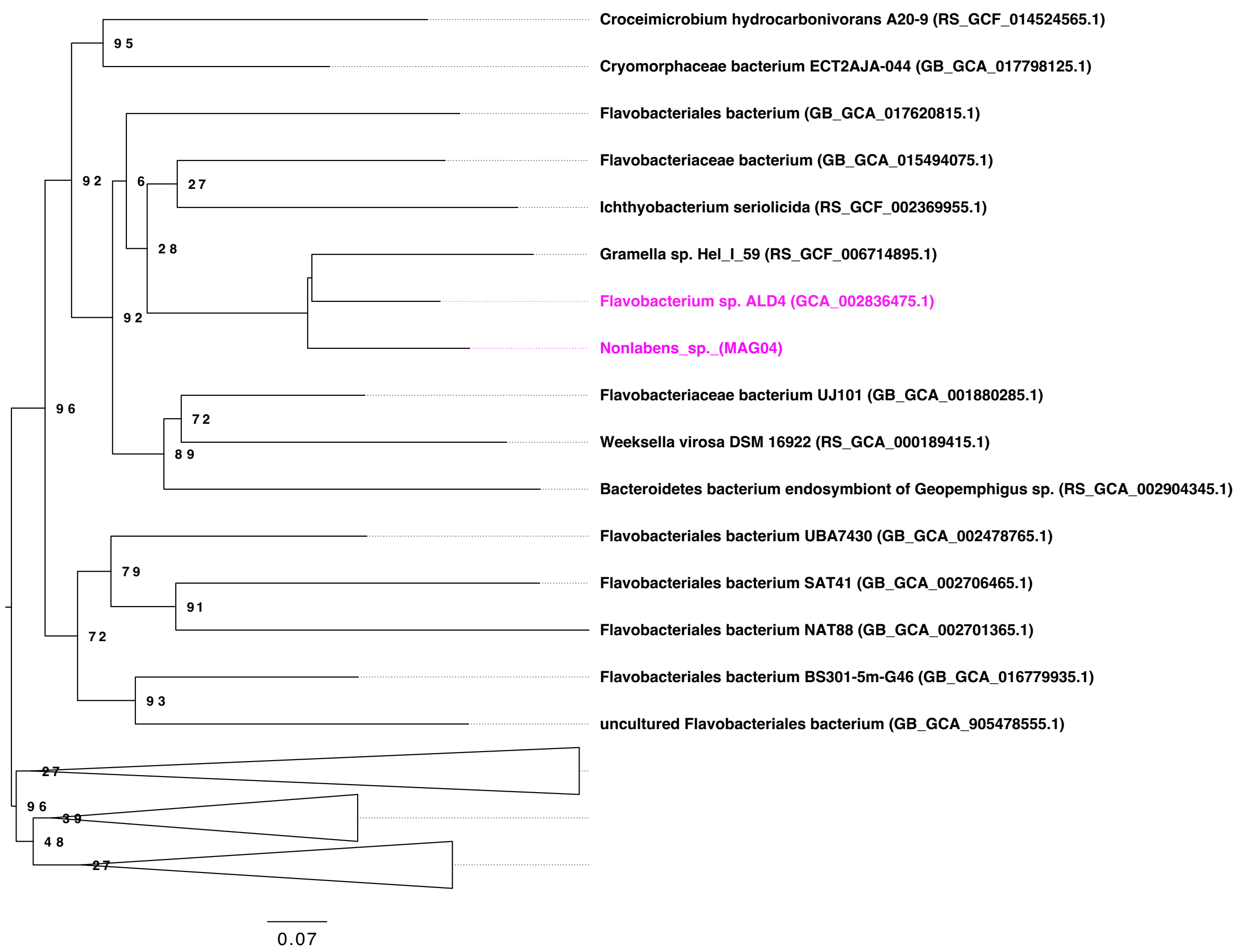


**Figure S1:** Tree showing phylogenetic relatedness genome "ext_mOTU_v3_31506" belonging to *Flavobacterium* sp. ALD4, to which melt pond sample reads mapped within the mOTUs tool and is related to *Nonlabens* sp. (MAG04) binned from the melt pond. The tree is derived from bac120.classify.tree (identification uses 120 bacterial marker genes) predicted by the classify_wf in GTDB-Tk v.2.1.0 [1] which uses pplacer v.1.1 [2] for the maximum-likelihood placement of genomes in the tree. A subnetwork of the full tree was extracted in Dendroscope v.3.8.8. [3] and the tree was rooted at the midpoint with the tips aligned in FigTree v.1.4.4 [4]. According to JSpeciesWS web server [5], ANIb for the two genomes is 67.31% with an aligned fraction of 28.05%.

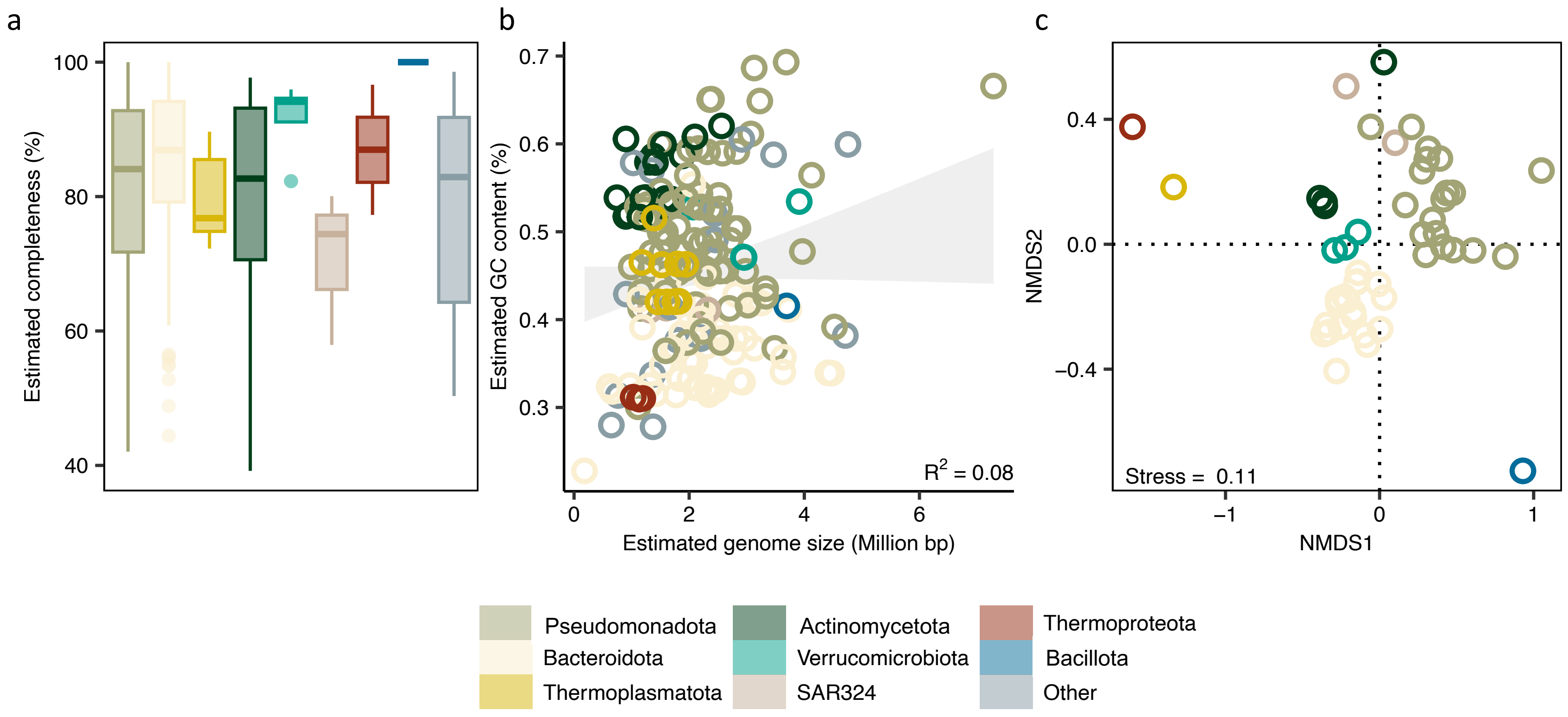


**Figure S2: Characteristics of the MAGs.** Estimated completeness grouped by phylum, based on CheckM2 [6] (a). Linear model of the estimated GC content (*y*) with the estimated genome size (*x*) revealing a low correlation ($R^2 = 0.02$, $p$-value = 0.34) (b). Divergence among MAGs based on functional orthologous genes (KO functional orthologs, accounting for multiple gene copies) using non-metric dimensional scaling (NMDS) (c).

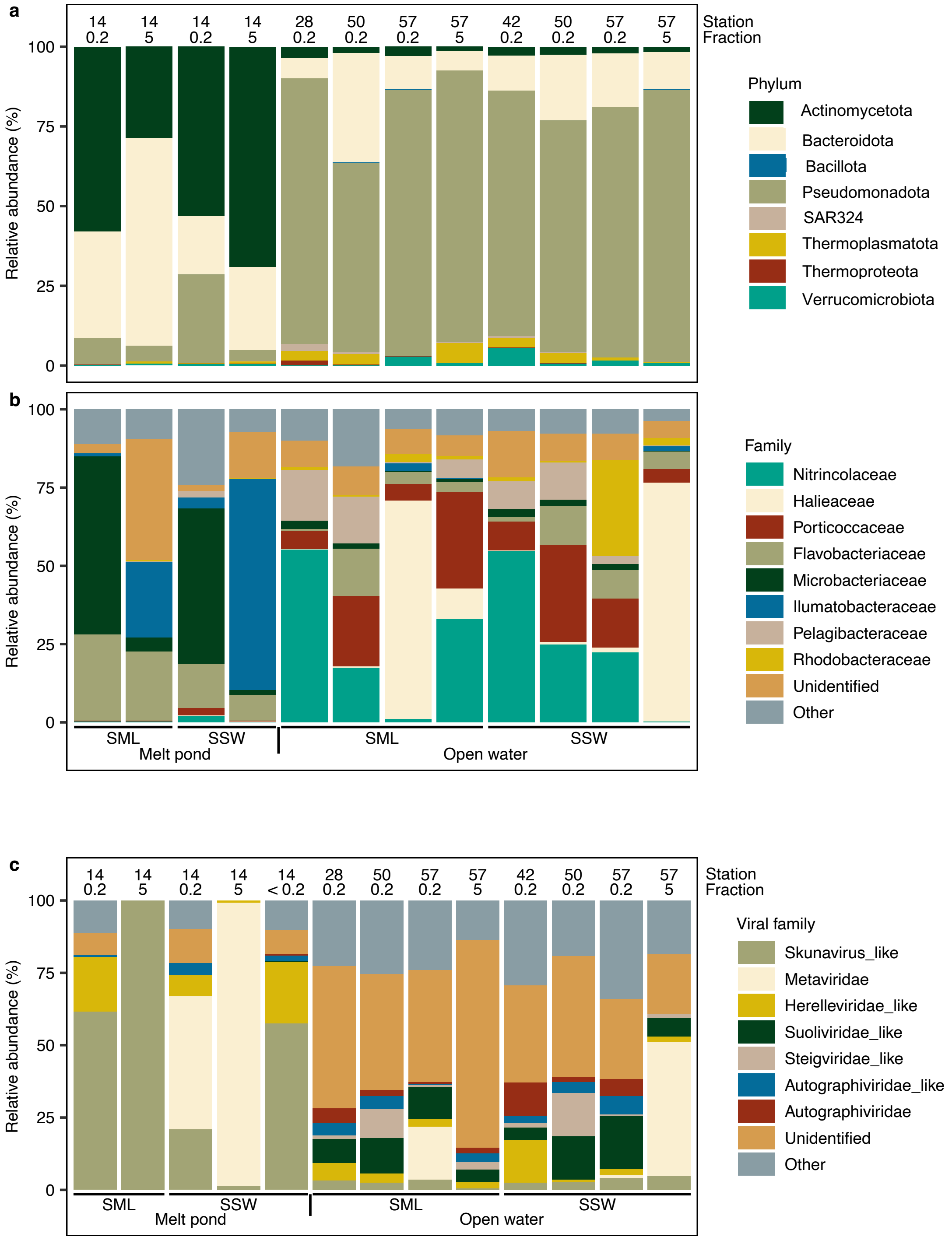


**Figure S3:** Taxonomic profiling expressed as relative abundance of the prokaryotic community based on metagenome assembled genomes (MAGs) at phylum (a) and family level (b). Relative abundances of viral families (c).

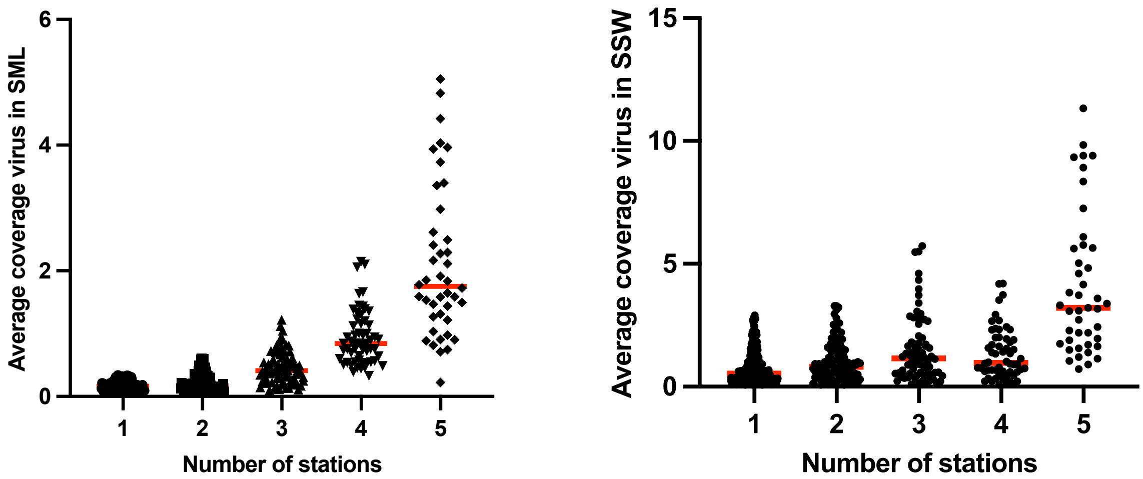


**Figure S4:** Correlation of the vOTU average coverage (red line indicates the median) for SML (a) and SSW (b) with number of stations a vOTU was present based on read breadth. This analysis shows that vOTU spread to different stations is more positively correlated to a higher vOTU coverage in the SML compared to SSW. The group comparison results are shown in the two tables below.

Dunn’s multiple comparison test results after Kruskal-Wallis test corresponding to SML groups from Figure S4 left plot

| Dunn's multiple comparisons test | Mean rank diff. | Significant? | Summary | Adjusted P Value |
|---|---|---|---|---|
| 1 vs. 2 | -27.08 | No | ns | 0.7891 |
| 1 vs. 3 | -114.9 | Yes | **** | <0.0001 |
| 1 vs. 4 | -196.5 | Yes | **** | <0.0001 |
| 1 vs. 5 | -240.9 | Yes | **** | <0.0001 |
| 2 vs. 3 | -87.85 | Yes | **** | <0.0001 |
| 2 vs. 4 | -169.5 | Yes | **** | <0.0001 |
| 2 vs. 5 | -213.8 | Yes | **** | <0.0001 |
| 3 vs. 4 | -81.61 | Yes | *** | 0.0002 |
| 3 vs. 5 | -126.0 | Yes | **** | <0.0001 |
| 4 vs. 5 | -44.35 | No | ns | 0.4346 |

| Test details | Mean rank 1 | Mean rank 2 | Mean rank diff. | n1 | n2 | Z |
|---|---|---|---|---|---|---|
| 1 vs. 2 | 100.7 | 127.8 | -27.08 | 107 | 92 | 1.757 |
| 1 vs. 3 | 100.7 | 215.6 | -114.9 | 107 | 76 | 7.068 |
| 1 vs. 4 | 100.7 | 297.2 | -196.5 | 107 | 58 | 11.12 |
| 1 vs. 5 | 100.7 | 341.6 | -240.9 | 107 | 42 | 12.20 |
| 2 vs. 3 | 127.8 | 215.6 | -87.85 | 92 | 76 | 5.229 |
| 2 vs. 4 | 127.8 | 297.2 | -169.5 | 92 | 58 | 9.324 |
| 2 vs. 5 | 127.8 | 341.6 | -213.8 | 92 | 42 | 10.59 |
| 3 vs. 4 | 215.6 | 297.2 | -81.61 | 76 | 58 | 4.318 |
| 3 vs. 5 | 215.6 | 341.6 | -126.0 | 76 | 42 | 6.043 |
| 4 vs. 5 | 297.2 | 341.6 | -44.35 | 58 | 42 | 2.019 |

Dunn’s multiple comparison test results after Kruskal-Wallis test corresponding to SSW groups from Figure S4 right plot.

| Dunn's multiple comparisons test | Mean rank diff. | Significant? | Summary | Adjusted P Value |
|---|---|---|---|---|
| 1 vs. 2 | -67.36 | Yes | ** | 0.0076 |
| 1 vs. 3 | -119.4 | Yes | **** | <0.0001 |
| 1 vs. 4 | -111.6 | Yes | *** | 0.0003 |
| 1 vs. 5 | -303.9 | Yes | **** | <0.0001 |
| 2 vs. 3 | -52.05 | No | ns | 0.6163 |
| 2 vs. 4 | -44.23 | No | ns | >0.9999 |
| 2 vs. 5 | -236.5 | Yes | **** | <0.0001 |
| 3 vs. 4 | 7.819 | No | ns | >0.9999 |
| 3 vs. 5 | -184.5 | Yes | **** | <0.0001 |
| 4 vs. 5 | -192.3 | Yes | **** | <0.0001 |

| Test details | Mean rank 1 | Mean rank 2 | Mean rank diff. | n1 | n2 | Z |
|---|---|---|---|---|---|---|
| 1 vs. 2 | 268.7 | 336.0 | -67.36 | 357 | 116 | 3.367 |
| 1 vs. 3 | 268.7 | 388.1 | -119.4 | 357 | 74 | 4.994 |
| 1 vs. 4 | 268.7 | 380.2 | -111.6 | 357 | 58 | 4.210 |
| 1 vs. 5 | 268.7 | 572.6 | -303.9 | 357 | 43 | 10.06 |
| 2 vs. 3 | 336.0 | 388.1 | -52.05 | 116 | 74 | 1.869 |
| 2 vs. 4 | 336.0 | 380.2 | -44.23 | 116 | 58 | 1.469 |
| 2 vs. 5 | 336.0 | 572.6 | -236.5 | 116 | 43 | 7.077 |
| 3 vs. 4 | 388.1 | 380.2 | 7.819 | 74 | 58 | 0.2382 |
| 3 vs. 5 | 388.1 | 572.6 | -184.5 | 74 | 43 | 5.140 |
| 4 vs. 5 | 380.2 | 572.6 | -192.3 | 58 | 43 | 5.105 |

a)

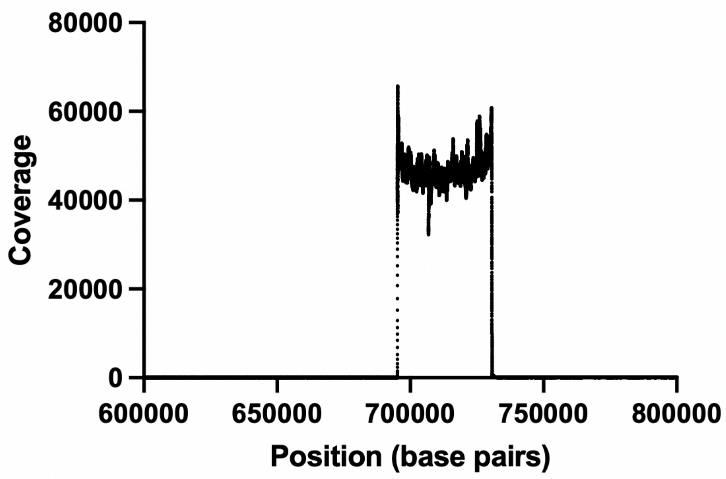


b)

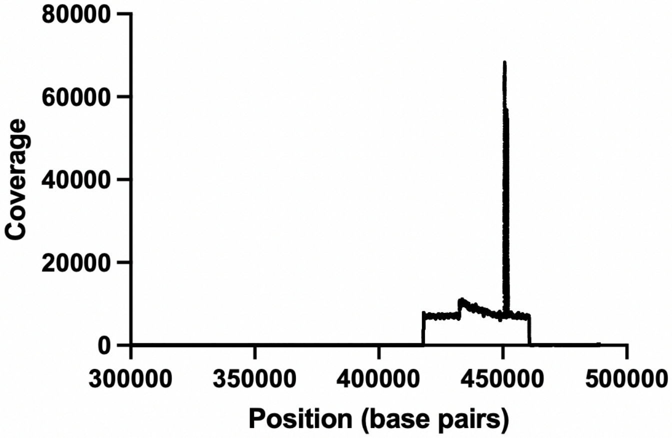


**Fig. S5:** Mappings of reads from sequenced DNA of the phage supernatant to the prophage carrying scaffolds of *L. aequorea* Arc30. Prophage 2, corresponding to Arctica_1 is located between position 694108 and 730554 bp (36.4 kb, a), and prophage 1, which corresponds to Arctus_1, between position 419014 and 463109 bp (44.1 kb, b).

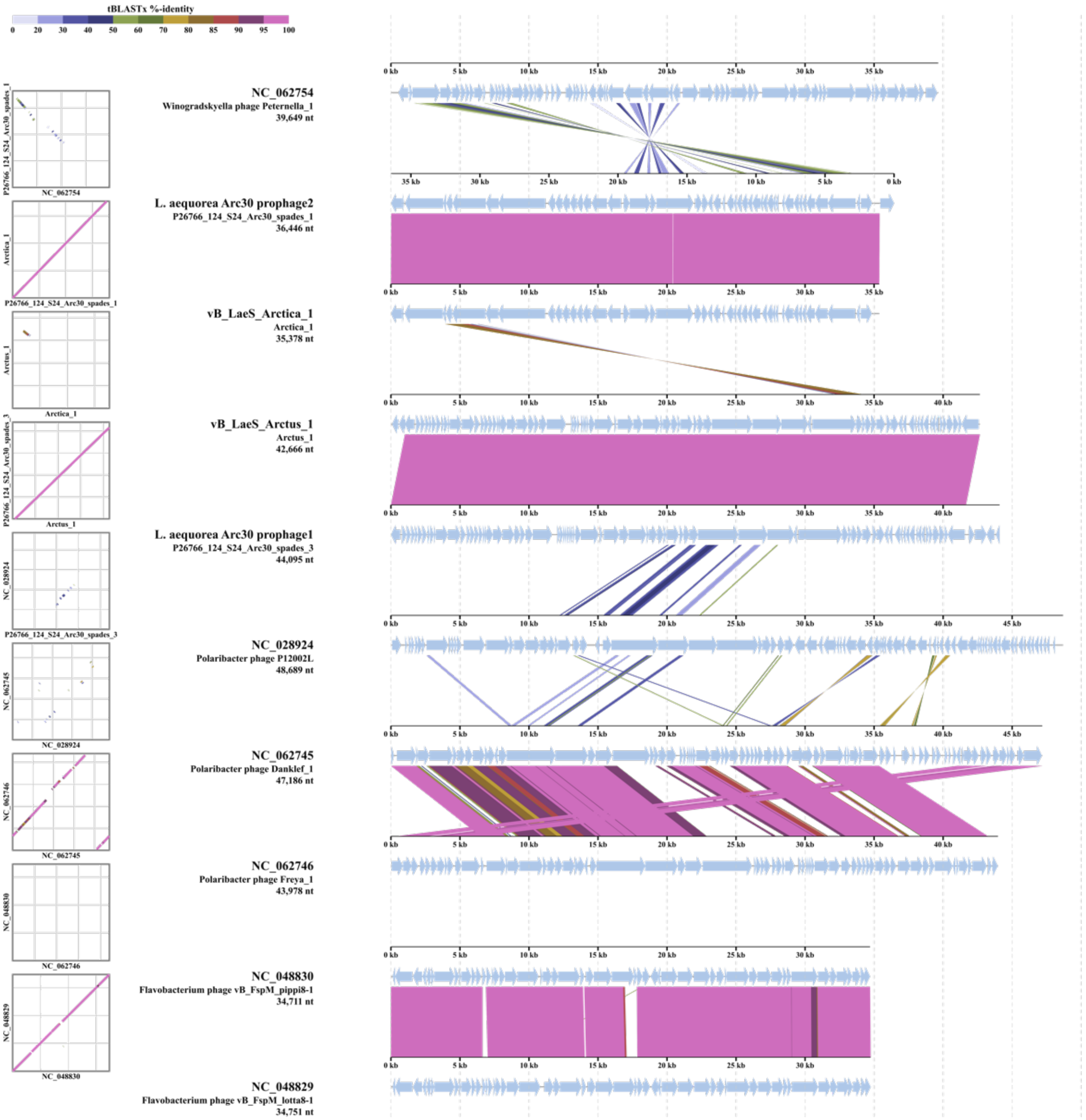


**Figure S6:** Alignments for vB_LaeS_Arctus_1 and vB_LaeS_Arctica_1 with related phages based on tBLASTx analysis conducted in VipTree v.4 [7].

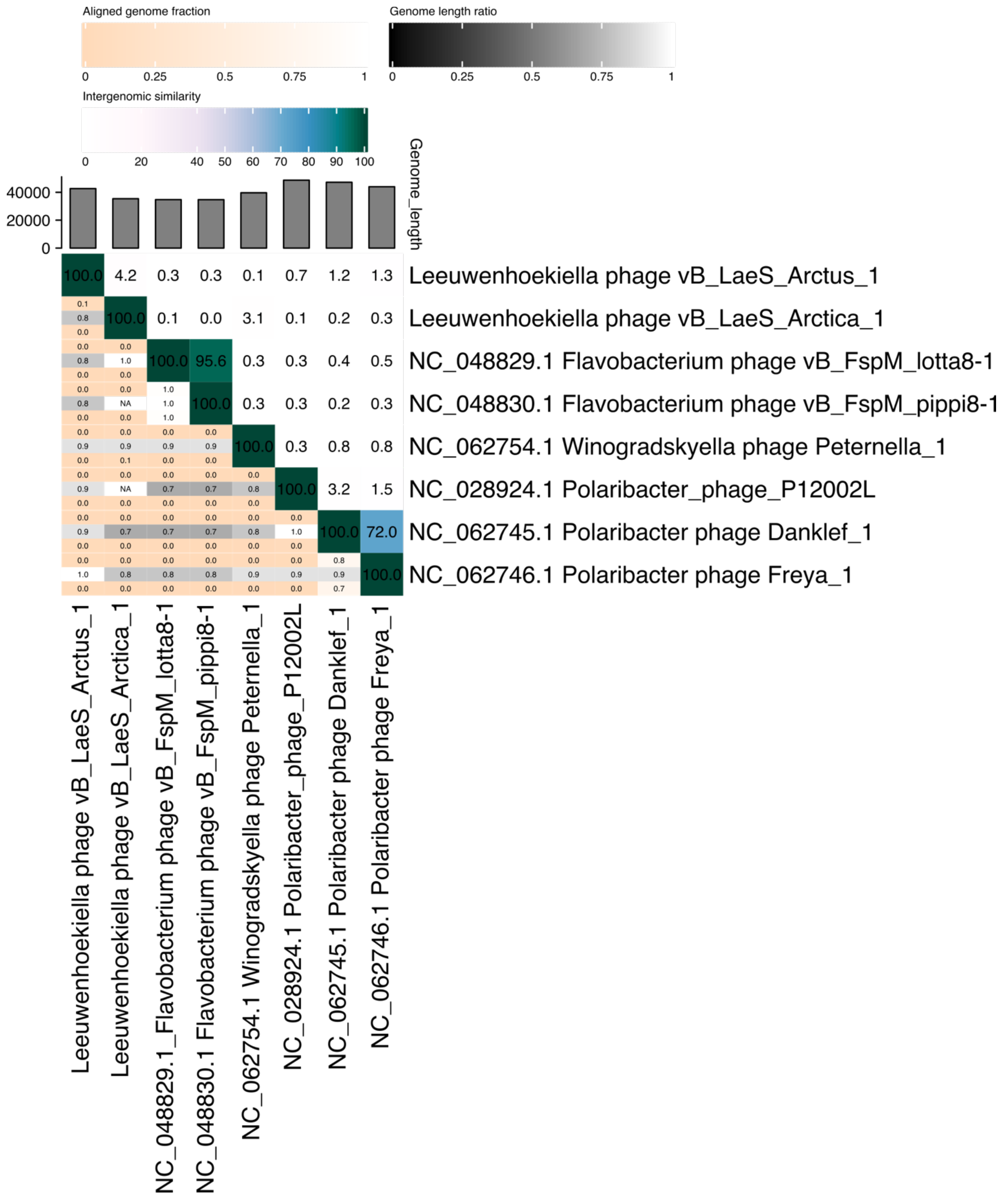


**Figure S7:** Heatmap derived from VIRIDIC [8] showing low intergenomic similarity between vB_LaeS_Arctus_1 and vB_LaeS_Arctica_1 and related phages as shown in Figure 7d.

**Issues with contaminations**

We were informed by the sequencing company about possible cross-contaminations in the sequencing plate as the plate's seal broke during shipment, which, after investigation, let us to avoid any inter sample comparisons (abundance, diversity, presence/absence, correlations) for the virome samples 118, 119, 121, 128.

Because we also had evidence for ship-borne water contamination in samples taken from icebreaker Oden's gangway, viruses that were assembled in samples 108, 111, 112, 113 and 128, were excluded from the sample pool unless they formed a species cluster with a vOTU from one of the other "clean" samples. Due to this issue, we also excluded bacterial MAGs from metabolic analysis if according to read breadth they were found at the gangway stations or are typical genera of non-marine, anaerobic bacteria like MAGs related to *Propionivibrio* sp., *Ruminococcus bromii, Sulfurospirillum* sp. All exclusions for MAG analysis have been further specified in Supplement Table S2 by labelling potential contaminants with a "C".

**>Arctus_1 Tape measure protein [PF20155.2] (db=pfam)**

MAKNQFQDAIEKAGLTLQEIEKRWISIDEKILQASKSASKLGKTDFNSAQPKDLNDRLQK
NATYRKQVNAEMKEQERLNKALAAAQAKFYSTQSGTNRQLQQTRFETNQLNAKYREQ
AILSSKLADEYQKQSTRLNMLRREAKAAAAQYGVNSKEAKNLIRDVNKLDASLKKVDA
AVGQHQRSVGNYGKAWQGVGKLMGAAVGAFGVYSAMQIGREIYAEIKAIDGLNKALK
QVTETTESYNQAKGFLGDLSQETGVQIKELTGAYLSFYAAAKNTNLTLEETQDIFRQTA
KAGATLGLSTEQVEGALRALEQMLSKGKVQAEEIRGQLGERLPGAFQILARSIGVSTAEL
DDMLKKGEVIADEVLPRFARELEKTFSLDKIDKVNTLAAAEGRLSTEWTRFVETLSNED
GAVTGFLTGTLELVTGIVSELRKLNEDLAPKSTSRVFDETIAKYKELGEAGKEEAMNQK
RNSEQAIKDAKNLQVILEERKNKLEEESGWFQNNFGKTKKEYKDLEDSIYNNNAKIAVQ
NGLLQAATEYLGLNTKEVEGNSKAEDENNDKKKKGIIALQGSIGAMEAVISKLEEEQSK
LATNGKEWSEYAGQINKAKDALQKIKTEYEGIEVLFEQEGVKDFDPFDYDLMDESQKR
ALKQVQDNAKAATNLLSDEYEKRLSTLEQFEKRKEDVLRDAARLEFDIRRDAFEKIADT
GQGFFQIEIDRYDQQIDALNENYDAQIEAAEGNEKQQQALRDEKLLKEQELERKKEEAE
KNAFLFSQGLALAQIGIDLARTISAIQVAATAMDALTPFAFGATGTTYRAANIPVAIGTA
AAQTALIAAQTIPALEQGDTTGKHEGQVMINDAKGAKFREIVQRTSGQMEVYSGRNVVI

DKKRGDKVYKAGQAPGGFDYNDLVNASINMSLADQYGRMSQAEAVQTFDFSAMESM
MDRKLSEFTKAVKTNKTVVNVPDSGASFAKAMRLNKIINK
